# Asgard archaeal origin of microtubules

**DOI:** 10.64898/2026.02.15.705738

**Authors:** Jan Löwe, Andriko von Kügelgen, Vicente J. Planelles-Herrero, Martin B. L. McAndrew, María A. Oliva, Julian Vosseberg, Stephan Köstlbacher, Jennah E. Dharamshi, Kathryn E. Appler, Fraser I. MacLeod, Stephanie-Jane Nobs, Steffen L. Jørgensen, Brendan P. Burns, Brett J. Baker, Tanmay A. M. Bharat, Emmanuel Derivery, Daniel Tamarit, Thijs J. G. Ettema

## Abstract

Eukaryotic cells change their shapes, actively segregate their DNA and contain membrane networks, facilitated by a complex cytoskeleton containing actin filaments, microtubules made from tubulin, and other components. These filaments have ancient evolutionary origins since actin- and tubulin-like proteins form prokaryotic cytoskeletons in archaea and bacteria. *Bona fide* eukaryotic F-actin can be traced back to crenarchaea and Asgard archaea, which are the closest known relatives of eukaryotes. A possible Asgard archaeal origin of microtubules was suggested recently with the discovery of a lokiarchaeon containing AtubAB mini microtubules that share architectural features with their eukaryotic counterparts. Using phylogenetic analyses of metagenomic data, here we report the broad occurrence of tubulins in Asgard archaea. Biochemical and structural analyses showed that one of our newly discovered heimdallarchaeial AtubAB tubulin pairs forms four-protofilament mini-microtubules that show dynamic instability and are inhibited by the tubulin drug maytansine. Our work raises the possibility that microtubule architecture and dynamics evolved in Asgard archaea prior to eukaryogenesis.

## INTRODUCTION

The eukaryotic cytoskeleton is one of the defining features of cellular complexity, enabling precise spatial organisation, cell division, directed transport, motility and the control of cell shape. It was shown previously that the two main filament-forming proteins of the eukaryotic cytoskeleton, actin and tubulin, have ancient prokaryotic (bacterial and archaeal) homologues, whose often dynamic filaments perform similar principal functions and have similar, but not identical structures (Löwe and Amos, 1998; van den Ent et al., 2001). Here, we studied the evolutionary history of one key component of the eukaryotic cytoskeleton, microtubules, which are made of tubulins.

The fundamental unit of microtubules consists of α and β tubulin heterodimers. These tubulins are closely related in amino acid sequence, each binding a GTP (guanosine triphosphate) nucleotide. The GTP bound to α tubulin, trapped in the middle of the heterodimer, is not hydrolysed because of K254 (pig tubulin numbering) in the T7 loop of β that protrudes into the GTP-binding site of α. The corresponding T7 loop of α that protrudes into the GTPase site of β has the catalytic E254 in the same position and facilitates polymerisation-dependent GTPase switching by completing the active site (Nogales et al., 1998a).

Heterodimers polymerise into alternating (… ABAB …) polar protofilaments (pfs) (Nogales et al., 1998b). In most eukaryotic microtubules, 13 pfs associate laterally to form a hollow cylindrical tube with a diameter of approximately 25 nm. Overall, 13-pf microtubules are a three-start pseudo-helix, with 12 times α-α and β-β lateral interactions and one seam that has α-β and β-α contacts (B and A lattices, respectively) (Zhang et al., 2015).

Microtubules can be highly dynamic, elongating and disassembling through the addition and loss of αβ tubulin heterodimers, a behaviour energetically driven by GTP binding, intrinsic GTPase activity upon polymerisation, and GDP to GTP exchange. Polymerisation triggers a conformational change, termed the cytomotive switch (Wagstaff et al., 2023), in both α and β tubulin, which also leads to a reorganisation of the α-β longitudinal interface. The cytomotive switch makes it possible for microtubule ends (and even single protofilaments) to have different rates of the addition of subunits, called the plus and minus ends.

Microtubules are most spectacularly utilised in the mitotic spindle of all eukaryotes, where their dynamics facilitate the search for chromosome attachments, and their depolymerisation drives chromosome segregation. One hallmark of microtubule dynamics is dynamic instability (Mitchison and Kirschner, 1984) that describes when heterodimer addition to the plus end is slower than GTP hydrolysis. This means the structure is no longer capped with stabilising GTP-bound tubulins and catastrophically depolymerises almost instantaneously, due to the strain the GDP state causes inside the microtubule.

Beyond α and β tubulins, additional members of the tubulin superfamily such as γ, δ, ε, and cryptic, carry out specific functions, including microtubule nucleation, basal body organisation, and the fine-tuning of microtubule architecture across diverse cellular contexts (Santana-Molina et al., 2023). The limited sequence similarity between eukaryotic tubulins and their known prokaryotic counterparts has long obscured the evolutionary origins of microtubules. Tubulin-like proteins (distant homologues of tubulin) in bacteria, archaea and viruses, such as FtsZ, CetZ, TubZ, PhuZ and artubulins (Wagstaff and Löwe, 2018) bear structural similarity to eukaryotic tubulins, form similar protofilaments but display only limited sequence similarity, and are typically separated by long branches in phylogenetic trees (Santana-Molina et al., 2023). They have not been reported to form microtubules.

In the early 2000s, members of the genus *Prosthecobacter* within the Verrucomicrobia were found to contain more closely related tubulin homologues, BtubA and BtubB (Jenkins et al., 2002), that form “mini microtubules” with similar properties to eukaryotic microtubules, but with greatly reduced protofilament numbers of 4-5 (Deng et al., 2017; Pilhofer et al., 2011). BtubAB bacterial tubulins, of which there are very few known sequences, cluster within the eukaryotic tubulin clade in phylogenetic trees (Santana-Molina et al., 2023), implying that they may have been acquired by horizontal gene transfer.

Research into the evolution of eukaryotes and their cytoskeleton, has recently been invigorated by the discovery of Asgard archaea [also known as Prometheoarchaeota (Imachi et al., 2024)], the closest known archaeal relatives of eukaryotes (Eme et al., 2023; Spang et al., 2015; Zaremba-Niedzwiedzka et al., 2017). Asgard archaeal genomes encode a diverse molecular toolkit of eukaryotic signature proteins (Vosseberg et al., 2024), with rapidly accumulating evidence indicating structural and functional conservation that predates eukaryogenesis, for example for actin (Akıl et al., 2021), endosomal sorting complexes required for transport (ESCRT) (Hatano et al., 2022) and soluble NSF attachment proteins (SNARE) (Neveu et al., 2020).

While close actin homologues can be found in most Asgard archaeal genomes (and crenactin in some crenarchaea, now classified as Thermoprotei) (Izoré et al., 2016), close tubulin homologues have only been found in Odinarchaeia (Zaremba-Niedzwiedzka et al., 2017) and in two related Lokiarchaeial species (Nobs et al., 2025; Wollweber et al., 2025). Both odinarchaeial and lokiarchaeial tubulins have been shown to form eukaryotic-like protofilaments (Akıl et al., 2022; Wollweber et al., 2025), with the lokiarchaeial tubulins AtubAB forming 5-pf mini microtubules (Wollweber et al., 2025). However, the unclear distribution of tubulin genes across Asgard archaeal genomes and the scarcity of detailed structural and functional studies currently limits our ability to infer their functional roles and to draw robust conclusions about the evolutionary origins of the eukaryotic microtubule cytoskeleton.

Here, we expand the set of known prokaryotic tubulins, with the aim of clarifying their evolutionary origin. We report the presence of close tubulin homologues in several novel Asgard archaeal metagenome-assembled genomes (MAGs), including in the closest Asgard archaeal relatives of eukaryotes. Phylogenetic analyses of these Asgard archaeal tubulin homologues revealed multiple tubulin paralogous copies clustering within the eukaryotic tubulin tree, revealing a complex evolutionary scenario of duplications and differential losses. Subsequent in-depth structural characterisation of Kariarchaeaceae (Heimdallarchaeia) AtubAB tubulin homologues revealed that their mini microtubules share many key characteristics of eukaryotic microtubules, including stable heterodimers, a seam, M-loops, maytansine inhibitor sensitivity, and dynamic instability - but differ uniquely in forming 4-pf microtubules.

## RESULTS

To establish the relationship between newly discovered prokaryotic sequences and eukaryotic tubulins, we performed sequence homology searches and phylogenetic analyses (Methods). These analyses revealed *bona fide* tubulin sequences in multiple prokaryotic groups (Figure 1A, Supplementary Figure S1A). The Odinarchaeial tubulins (hereafter “Odin tubulins”) cluster as a monophyletic group, formed by proteins encoded in a single copy in all known Odinarchaeial genomes, indicative of a conserved function ancestral to this group. We identified a single Bathyarchaeia tubulin sequence clustered through a long branch, indicative of an ancient horizontal gene transfer (HGT), but contamination of this sequence could not be ruled out. Novel Verrucomicrobia sequences formed monophyletic groups with the BtubA and BtubB clades, expanding the known sequence diversity for these bacterial tubulins that were previously shown to form 4- or 5-pf (protofilament) mini microtubules (Deng et al., 2017; Pilhofer et al., 2011).

**Figure 1.**
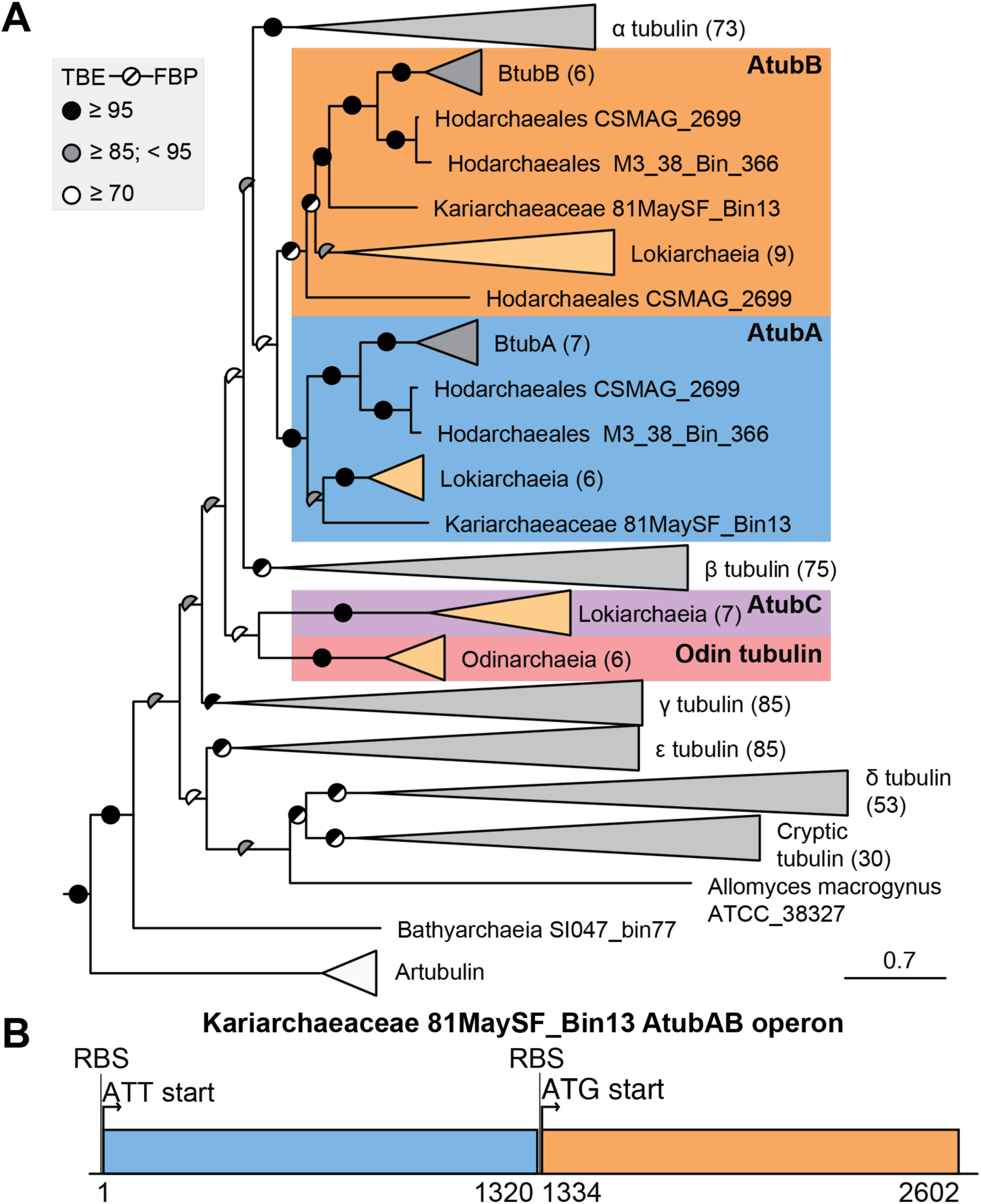
Phylogeny of tubulins reveals Asgard archaeal AtubAB tubulin genes. **A)** Colours for collapsed clades indicate taxonomy: Asgard archaea (orange), eukaryotes (grey), bacteria (dark grey) and other archaea (white). Branch support values represent Transfer Bootstrap Expectation (TBE) and standard Felsenstein’s Bootstrap Support (FBS). Phylogeny reconstructed using IQ-TREE 3 under the model Q.pfam+C50+G4+PMSF. The outgroup was removed for visualisation. Branch length scale indicates substitutions per site. The full tree is shown in Supplementary Figure S1A, gene maps in S1B. The alignment (trimmed off sites with over 90% gaps and sequences with over 50% gaps) contains 477 sequences and 578 columns. **B)** Schematic of the tubulin-like Kariarchaeaceae 81MaySF_Bin13 AtubAB operon, coding for the Kari AtubAB proteins used in this study (see also Supplementary Figures S1C & D). Arrows indicate start codons. RBS, ribosome binding site.

Additional tubulin copies were found in Loki- and Heimdallarchaeia. One of these, encoded in the genome of *Candidatus* Lokiarchaeum ossiferum, has been shown recently to form mini microtubules (Wollweber et al., 2025) that recapitulate many structural features of eukaryotic 13-pf microtubules, but not protofilament number. We found multiple additional tubulin homologs in Lokiarchaeia, including MAGs recovered from marine sediments sampled in the Arctic ocean, the Gulf of Mexico, and the Australian Shark Bay. Additionally, three heimdallarchaeial MAGs were also found to contain tubulins. The lokiarchaeial, odinarchaeial and heimdallarchaeial tubulins formed three different monophyletic groups, which we named AtubA, AtubB and AtubC (Figure 1A). AtubA and AtubB were found to also contain BtubA and BtubB, respectively, which clustered with the heimdallarchaeial tubulins with high confidence, indicating an Asgard archaeal origin of the verrucomicrobial tubulins. As bacterial tubulins were previously thought to have evolved from eukaryotic tubulins (Schlieper et al., 2005), these results shed new light on a long-standing debate in tubulin evolution. The AtubC clade comprises tubulin sequences from Lokiarchaeia.

AtubA, AtubB and AtubC cluster confidently with the eukaryotic tubulin clade in most phylogenetic reconstructions (see Supplementary Discussion). In those, AtubA and AtubB mostly cluster as a sister group to α tubulin. The AtubC clade has a more uncertain position yet is placed at the base of the α and β tubulin clade in most analyses, albeit with low support. Odin tubulins tend to cluster together with AtubC, indicating they may be orthologous to the Lokiarchaeial AtubC sequences (Supplementary Discussion). The position of artubulins, linked through a very long branch to the eukaryotic tubulin clade, is never fully resolved, with main placements being at the base of the eukaryotic tubulin group or within, possibly as sister to Odin tubulins (Supplementary Discussion).

We found AtubA and AtubB to always be encoded by genes located in tandem, and AtubC is encoded within 10 kb of these genes in Lokiarchaeia (Supplementary Figure S1B). A second AtubB homolog (AtubB2) is found in some lokiarchaeial genomes, at a distant position. The genomic context of these genes seems not to be conserved between taxonomic groups.

To further our understanding of the evolutionary and functional relationships we set out to investigate the structure and dynamics of the newly discovered Asgard Atub tubulins. One such AtubAB pair turned out to be highly accessible biochemically: Kariarchaeaceae 81MaySF_Bin13 AtubAB, hereafter AtubAB (Figure 1B, Supplementary Figure S1C). AtubA and B are encoded in an operon, with the second ribosome binding site (RBS) for *atubB* partly overlapping with the stop codon of *atubA* (Figure 1B). AlphaFold 3 predictions (Abramson et al., 2024) revealed AtubAB to likely have the tubulin fold, form heterodimers and to form alternating (ABAB) protofilaments (Supplementary Figure S1D). Inspection of the T7 loops (Nogales et al., 1998a) in the predicted protofilaments also revealed that it is likely that the GTPase site between AtubA at the top and AtubB at the bottom in the dimer is not active because of AtubA K248 not supporting hydrolysis in the GTP-binding pocket of BtubB below. In our hands, AlphaFold 3 could not predict the correct filament structure of AtubAB.

For bacterial expression of AtubAB we constructed two bicistronic expression vectors, pHis17-AtubAB for untagged expression and pHis17-AtubAhBs for His_8_(h)- and Strep(s)-tagged AtubAB, respectively (Methods, Supplementary Figures S2A & B). Small-scale expression tests revealed that AtubAB express highly in *Escherichia coli* C41(DE3) cells, making up a large proportion of the soluble protein content of the cells (Figure 2A, left). Nickel-based purification of AtubAhBs pulled out AtubAh and AtubBs simultaneously (Figure 2A, middle), even though only AtubAh had a His_8_ tag, indicating that AtubAB form a stable heterodimer. Large amounts of protein (>100 mg) could be purified, which led us to attempt purifying untagged AtubAB by anion exchange chromatography, followed by size exclusion chromatography (SEC, Methods). Again, large amounts of protein could be purified (>100 mg) and AtubAB co-fractionated throughout the procedure, despite not adding GTP to the buffers and using high salt conditions during the anion exchange chromatography (Figure 2A, right). SEC coupled with multi-angle light scattering (SEC-MALS) of the purified untagged AtubAB sample revealed a single peak with an estimated mass of 91 kDa, close to the expected mass of 94 kDa of the AtubAB heterodimer (Figure 2B). AtubAB is a slow GTPase under the conditions used: 0.192 +/- 0.004 nmol phosphates released/nmol AtubAB/min (25°C; Figure 2C, Methods).

**Figure 2.**
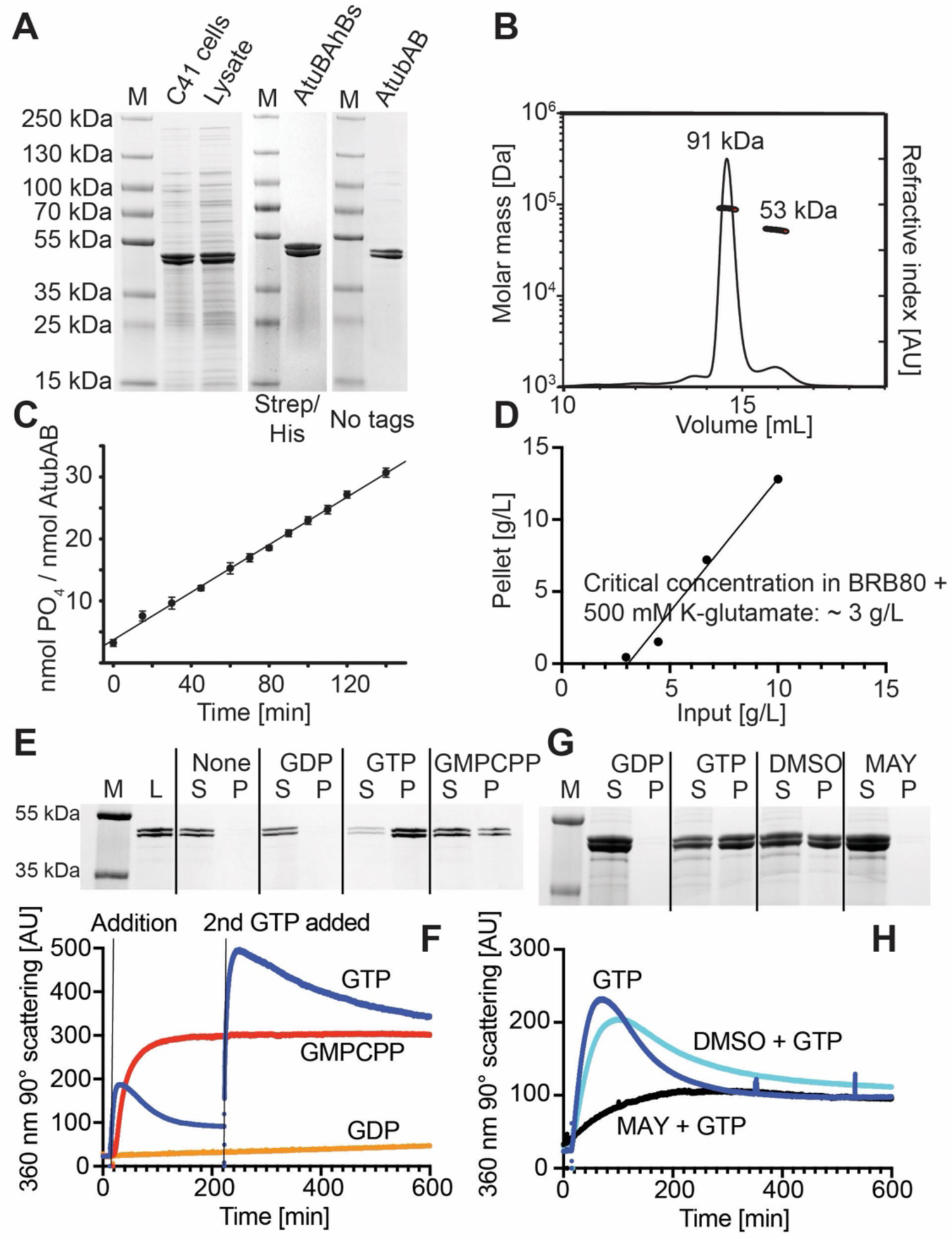
AtubAB expressed in *E. coli* forms stable heterodimers and polymerises in the presence of GTP, which is inhibited by the tubulin inhibitor maytansine. **A)** Coomassie-stained SDS PAGE gels showing strong soluble untagged AtubAB expression in *E. coli* C41(DE3) cells (left), purified His_8_ and Strep-tagged AtubAhBs (middle) and purified untagged AtubAB (right), as used in this study. **B)** Size-exclusion chromatography coupled with multi-angle light scattering (SEC-MALS) reveals that purified untagged AtubAB is a stable heterodimer. **C)** AtubAB is a slow GTPase: 0.192 +/- 0.004 nmol phosphates released/nmol AtubAB/min (25°C), *n* = 3. **D)** Critical concentration of AtubAB polymerisation in BRB80 + 500 mM potassium glutamate Polymerisation Buffer (used throughout this study) is ∼3 g/L (62 µM dimer). **E)** Pelleting assay revealing that AtubAB polymerisation is GTP-dependent and can also be triggered by the non-hydrolysable GTP analogue GMPCPP. L, loaded; S, supernatant; P. pellet. **F)** 90° 360 nm light scattering assay showing the same GTP/GMPCPP-dependence of AtubAB polymerisation as in E. GTP hydrolysis leads to depolymerisation, adding GTP again leads to re-polymerisation. **G)** Pelleting assay showing the inhibition of GTP-induced AtubAB polymerisation by the tubulin inhibitor maytansine (MAY). DMSO and MAY reactions also contained GTP. (see also Supplementary Figure S2D) **H)** 90° 360 nm light scattering assay showing the same maytansine inhibition of GTP-induced AtubAB polymerisation as in G.

Attempts to polymerise AtubAB revealed that standard tubulin buffers such as BRB80 (Methods) did not yield filaments as investigated by pelleting and negative staining electron microscopy. After screening other conditions, we settled on the inclusion of 500 mM potassium glutamate in BRB80 as the Polymerisation Buffer used throughout this study. This has been used early with eukaryotic tubulin (Hamel and Lin, 1981) and was also found to polymerise Lokiarchaeial AtubAB best (Wollweber et al., 2025). The critical concentration of untagged AtubAB in Polymerisation Buffer in the presence of GTP was determined by a pelleting assay to be ∼3 g/L (32 µM dimer) (Figure 2D). Pelleting revealed that the polymerisation of AtubAB is strictly dependent on GTP or its non-hydrolysable analogue, GMPCPP (guanylyl 5’-α,β-methylenediphosphonate; Figure 2E). It also showed that AtubAB pellet together, stoichiometrically. Ninety degrees 360 nm light scattering was used as an alternative method to confirm the nucleotide dependence of polymerisation of AtubAB, and to show GTP hydrolysis-driven depolymerisation, which was absent in the GMPCPP-polymerised sample (Figure 2F). When we analysed the dynamics of tagged AtubAhBs protein in the same light scattering assay, we found that the tag altered its behaviour, leading to enhanced polymerisation and faster depolymerisation (Supplementary Figure S2C).

Since the AtubAB proteins are closely related to eukaryotic tubulins in sequence (∼35-43% sequence identity, depending on the tubulins used), we then tested an array of tubulin-directed drugs and compounds that either stabilise or disrupt microtubule formation (Supplementary Figure S2D). Maytansine (MAY), a small molecule compound preventing microtubule polymerisation by binding and sequestering tubulin dimers, showed strong inhibition of AtubAB polymerisation, both in pelleting and in the light scattering assay (Figures 2G & H). A binding assay with a fluorescein-maytansinoid (FcMaytansine) revealed a K_d_ of 87 ± 17 nM (mean ± SEM, *n* = 4, see Methods, Figure S2Ei), to be compared with 6.8 nM for FcMaytansine:tubulin (Menchon et al., 2018). A subsequent fluorescence anisotropy competition experiment with FcMaytansine and maytansine revealed a K_d_ value for (unlabelled) maytansine against AtubAB of 5.1 ± 0.2 µM. This is in the same range as maytansine binding to eukaryotic tubulin [∼900 nM, (Lopus et al., 2010)] (Figure S2Eii). A superposition of the AtubAB AlphaFold 3 model with a structure of maytansine bound to β tubulin (PDB 4TV8) (Prota et al., 2014) revealed that the maytansine binding site is highly conserved (Supplementary Figure S2F). We conclude that AtubAB is similar to eukaryotic αβ tubulins in terms of its biochemical properties, polymerisation behaviour and its susceptibility to the inhibitor maytansine.

To determine the structure of AtubAB was challenging because the organism from which the AtubAB metagenomic sequences came originally has not been cultivated or isolated. Therefore, we first determined the in-cell structure of AtubAB after over-expression in *E. coli* C41(DE3) cells as described above (Figure 2A, left). The cells were thinned by focussed ion beam (FIB)-milling into lamellae <200 nm thick (Supplementary Figure S3A). Subsequent electron cryotomography (cryo-ET) of the lamellae revealed bundles of filaments filling large areas of the *E. coli* cells in a way that could interfere with cell division (Figure 3A, Movie M1). Some single filaments were identified. These mostly appeared as two lines of density along the filament axis (Figure 3A, right). Model-free subtomogram averaging of the filaments using helical symmetry as implemented in Relion 5 (Burt et al., 2024) (Methods, Supplementary Figure S3B) resulted in a 6.4 Å-resolution map that revealed a 4-stranded (4-pf), slightly twisting mini microtubule structure (Figure 3B). Symmetry parameters were twist = -89.9°, rise = 10.3 Å.

**Figure 3.**
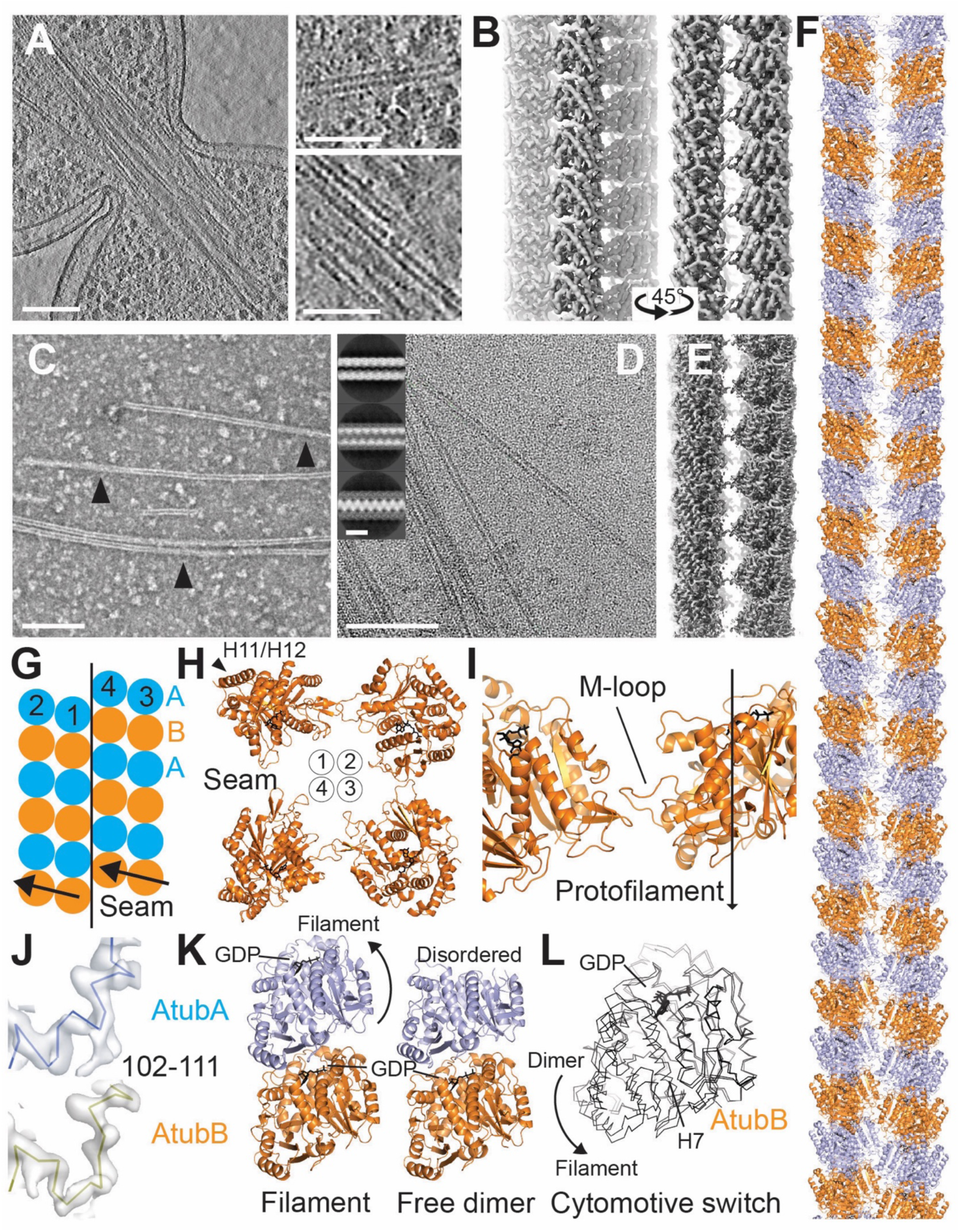
AtubAB cryo-EM mini microtubule structures in-cell in *E. coli* after over expression, and *in vitro* after purification. **A)** Electron cryotomogram of a focussed ion beam (FIB)-milled lamella of an *E. coli* C41(DE3) cell overexpressing untagged AtubAB, revealing bundles of filaments passing a cell division site (left overview, right magnified views). See also Movie M1. Scale bars 100 nm (left), 50 nm (right). **B)** Model-free helical subtomogram average (Supplementary Figure S3B), calculated from tomograms as in panel A, revealing a four-stranded mini microtubule at 6.4 Å resolution. **C)** Negative staining electron micrograph of purified, untagged AtubAB polymerised in BRB80 + 500 mM potassium glutamate Polymerisation Buffer, revealing filaments as seen in A and B (scale bar 100 nm). The arrowheads mark crossovers, indicating slowly twisting filaments. **D)** Cryo-EM micrographs of AtubAB polymers in Polymerisation Buffer (scale bar 100 nm); insets: 2D classes after helical segment picking and image processing (scale bar 10 nm). **E)** Cryo-EM structure of purified untagged AtubAB filaments after helical single particle processing at a resolution of 3.1 Å, revealing the same four-stranded mini microtubules as in panel B (Supplementary Figure S3E). **F)** Fitted and refined atomic model of the AtubAB four-stranded mini microtubule. AtubA blue, AtubB orange. See also Movies M2 & M3. **G)** Schematic of the pseudo-helical symmetry of AtubAB mini microtubules, leading to a seam. **H)** Top view of the four AtubAB mini microtubule protofilaments. Helices H11/H12 of the tubulin fold point outwards. **I)** AtubAB mini microtubule protofilaments are held together by M-loops (residues ∼270-280). See also Movie M3. **J)** Cryo-EM density regions demonstrating the identification of AtubA and AtubB after symmetry expansion and 3D classification. **K)** Comparison of the AtubAB heterodimer in the mini microtubule filament (as shown in panel F, left) and the structure of un-polymerised, untagged AtubAB as determined by single particle cryo-EM (right). **L)** Superimposing the AtubBs in panel K reveals the polymerisation-dependent cytomotive switch in AtubAB. See also Movie M4.

To reveal further molecular details of the AtubAB mini microtubules, we turned to electron microscopy (EM) of purified material, using untagged AtubAB protein in the BRB80 Polymerisation Buffer containing 500 mM potassium glutamate. Negative staining electron microscopy revealed slowly twisting filaments with two parallel lines and occasional crossovers (Figure 3C). Filament formation was again found to be strictly dependent on the presence of GTP or GMPCPP (Supplementary Figure S3C). In electron cryomicroscopy (cryo-EM), the filaments were often bundled, so manual picking was employed to pick single filaments (Figure 3D). Subsequent particle extraction and 2D classification revealed classes that are consistent with a 4-pf structure (Figure 3D, insets). Helical reconstruction of the cryo-EM data produced a map at 3.1 Å resolution that was very similar to the cryo-ET in-cell structure, but at higher resolution. The reconstruction used helical symmetry with a twist of - 89.84° and a rise of 10.46 Å (Figure 3E, Supplementary Figures S3D & E). Since AtubAB formed stable heterodimers even outside of the filaments (Figure 2B) and pelleted filaments contained both proteins (Figure 2E), we assumed that the structure consists of ABAB alternating protofilaments (Figure 3F). As a consequence, the helical symmetry observed means that the structure must be pseudo-helical and contain a seam, just as in eukaryotic 13-pf microtubules, (Figures 3G & 3H), in which lateral contacts change from A-to-A and B-to-B, to A-to-B and B-to-A (called B and A lattices, respectively, in eukaryotic microtubules). Moreover, the C-terminal helices H11 and H12 of the tubulin fold point outwards in AtubAB mini microtubules (Figure 3H), just as they do in microtubules.

The way protofilaments in AtubAB mini microtubules are held together laterally is highly reminiscent of eukaryotic microtubules (Figure 3I). An M-loop (microtubule loop) (Nogales et al., 1999), formed in AtubA and AtubB by residues ∼270-280, reaches over from the C-terminal domain of one subunit to the GTPase domain of another, contacting a hydrophobic pocket mainly through a single phenylalanine residue (F275 in AtubA and F278 in AtubB). The M-loop is the only inter-protofilament contact holding the entire structure together through repeated contacts that increase the avidity of this interaction.

We then aimed to differentiate AtubA and AtubB in the cryo-EM map. For this we used 8-fold symmetry expansion with subsequent 3D classification, focussing on a single dimer (Methods, Supplementary Figure S3D, box). The resulting map allowed AtubA and B to be distinguished both visually in a specific region of the map (Figure 3J) and computationally, by automated model building using ModelAngelo (Jamali et al., 2024), which produced complete atomic models for both AtubA and AtubB (Supplementary Figure S3D). While our procedure did not resolve the seam position in each mini microtubule observed (for which there was not enough signal because of the 38% sequence identity between AtubA and B), it was sufficient to reveal the alternating nature of each protofilament and enabling us to build a complete AtubAB mini microtubule structure with a seam, as required by the symmetry (Figures 3F & 3G, Movie M3).

Comparison of the resulting structure with the cryo-ET in-cell structure revealed an excellent fit, meaning that AtubAB forms the same 4-pf mini microtubules when expressed in *E. coli* and when polymerised *in vitro* (Supplementary Figure S3E). Comparison of the AtubAB heterodimer in the filament with eukaryotic αβ tubulin revealed very small deviations, with RMSDs between AtubA and β tubulin (PDB 7QUP) of 0.9 Å and between AtubB and α tubulin PDB 7QUP) of 0.92 Å, emphasising their close similarity at the subunit level (Supplementary Figure S3F, Movie M2). Surprisingly, we found that the AtubAB mini microtubules only contain GDP as indicated by the map. This matched our biochemical finding that purified AtubAB contained only GDP (Supplementary Figure S3G). The lack of un-hydrolysed GTP trapped between AtubA and B, as is seen in tubulin between α and β tubulin is likely due to the time taken for purification using anion exchange and SEC, which took a day and was performed in buffers that lack GTP (Methods). Interestingly, the AtubAB GDP heterodimer was still stable (Figure 2B).

Since dynamic actin and tubulin filaments undergo a cytomotive switch (Wagstaff et al., 2023) that depends on polymerisation-induced changes in monomer structure, we compared monomer and polymer structures of AtubAB. Using a cryo-EM single particle analysis (SPA) we solved the structure of un-polymerised and untagged AtubAB at 3.5 Å resolution (Supplementary Figures S3H & I). Parts of the GTPase site on top of AtubA were not resolved in the map, likely because no nucleotide had been added to the sample, meaning the site might be empty and hence disordered (Figure 3K). This is likely due to our purification protocol being unable to maintain nucleotide binding in the solvent-exposed AtubA GTP binding site. Comparison of the AtubAB structures in the filament and un-polymerised (free) heterodimer forms revealed two major differences. First, the inter-subunit angle changed. Second, each subunit underwent tubulin’s canonical polymerisation-state-dependent cytomotive switch, in which the C-terminal domain, including Helix 7 (H7) (Nogales et al., 1998a), moves downwards upon polymerisation (Figures 3K & L, Movie M4). This is the same conformation change as in all other tubulins and tubulin-like proteins (Wagstaff et al., 2023). Data of the structural work are summarised in Supplementary Table 1.

The structural work revealed that AtubAB forms mini microtubules (Movie M3) that share many features with eukaryotic microtubules, including: the subunit fold (Movie M2), overall polarity, alternating protofilaments, polymerisation-dependent GTPase activity, pseudo-helical symmetry with a seam, C-terminal H11/H12 helices on the outside, very little overall twist, M-loops holding protofilaments together and, importantly, the polymerisation-dependent cytomotive switch.

Having established strong biochemical and architectural similarities of AtubAB 4-pf mini microtubules with eukaryotic 13-pf microtubules, we set out to investigate filament dynamics. We labelled GMPCPP-bound stable AtubAB filament seeds with Alexa Fluor 647 and GTP-bound AtubAB with Atto 488. Some AtubAB was biotin-labelled to constrain filaments to the biotin-functionalised glass surface with neutravidin (Figure 4A, Methods). Total internal reflection fluorescence (TIRF) microscopy was then used to image the filaments, seeded by the red GMPCPP seeds, as they grew and shrank. The filaments exhibited two different growth speeds (Supplementary Figures S4A & B, Movie M5) (minus ends, median speed of growth 0.48 µm/min; plus ends, median speed of growth 1.80 µm/min), which is caused by their structural polarity and the kinetic difference of subunit addition to either end as caused by the cytomotive switch (Wagstaff et al., 2023).

**Figure 4.**
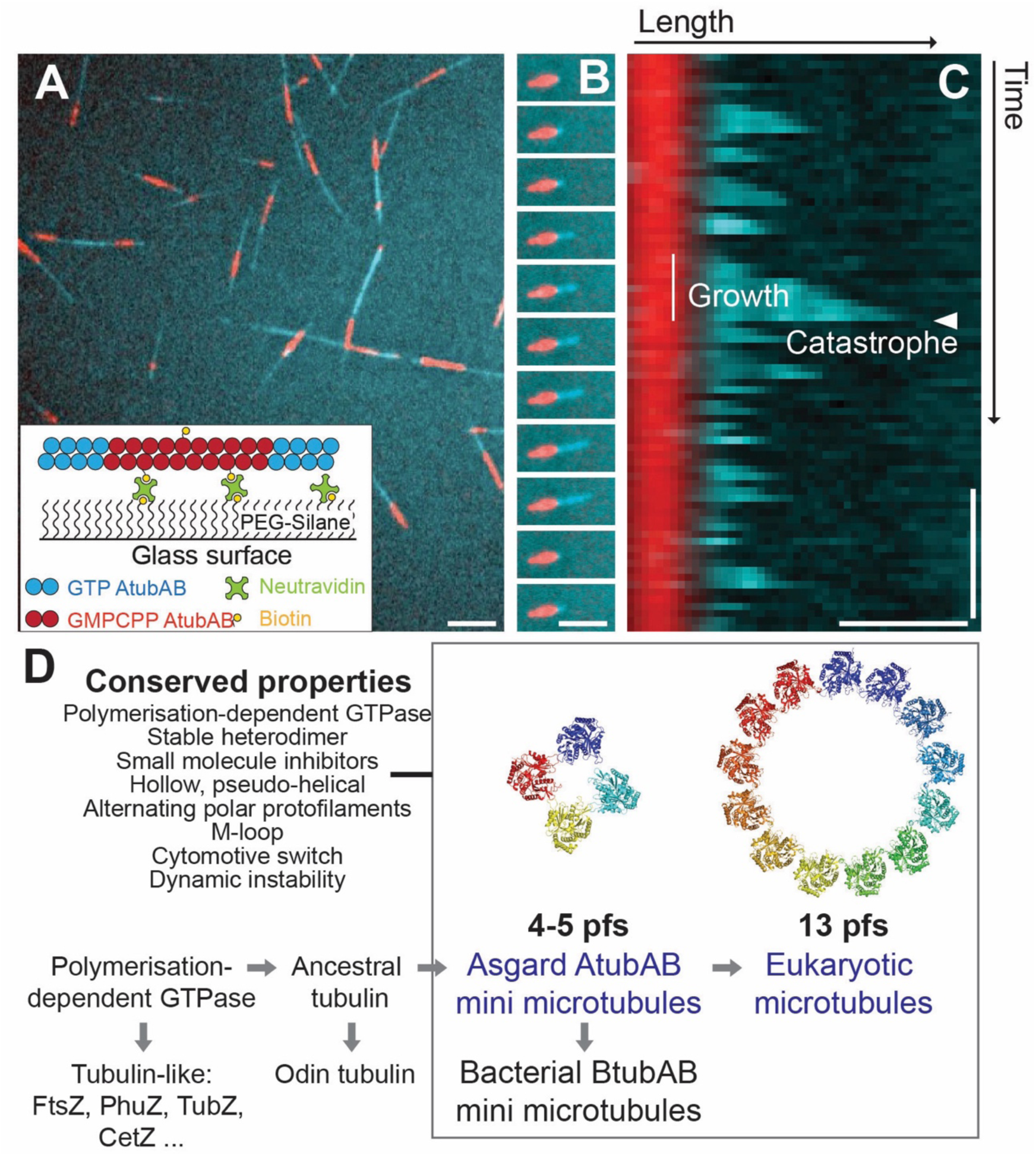
AtubAB dynamic instability revealed by TIRF microscopy. **(A)** Inset: assay design; biotinylated, GMPCPP-stabilised AtubAB dimers (labelled with 20% Alexa Fluor 647, shown as red) were anchored via NeutrAvidin to PEG Silane. Free AtubAB dimers (4 µM 20% Atto 488-labelled, cyan) were added, and filament polymerisation was observed by TIRF (total internal reflection fluorescence) microscopy. Field view revealing red GMPCPP seeds with plus- and minus-end growth (cyan). Only this used 20% tagged (unlabelled) AtubAhBs to observe dynamics at high concentrations needed for field views. Scale bar = 10 µm. **B)** Time-lapse of plus-end growth of untagged AtubAB filament. The 10^th^ image down shows the moment just after catastrophe, the ultra-rapid depolymerisation of the mini microtubule. Scale bar 5 µm. **C)** Kymograph (time vertical, filament axis horizontal) showing an untagged AtubAB filament growing and undergoing catastrophe at the plus-end. One growth episode (line) and one catastrophe event (arrowhead) are highlighted - it is the same as in panel B. Scale bars, 2 min / 2 µm. See also Supplementary Figures S4A & B and Movie M5. **D)** Asgard archaeal 4-pf mini microtubules show all the hallmarks of eukaryotic 13-protofilament microtubules - but are much thinner. We therefore suggest it is possible that Asgard mini microtubules are the evolutionary precursors of eukaryotic microtubules. Arrows do not indicate evolutionary distances or complexity.

The fast growing plus end showed strong dynamic instability (Mitchison and Kirschner, 1984) (Figures 4B & C, Supplementary Figures S4A & B). After a period of plus end growth, the filaments underwent a change in behaviour, leading to a process called catastrophe that involves rapid depolymerisation. The polymer, which is then thought to be completely in the GDP-bound form, loses its protection through a cap of GTP-bound subunits, which allows internal strain to be released via rapid polymer disassembly. Supplementary Figure S4A shows two more examples of filaments exhibiting dynamic instability at the plus end, growing to the right. They also show the slower minus end growth (to the left), which does show dynamic instability under the conditions used. Supplementary Figure S4A, iv shows the effect of adding in tagged AtubAhBs, which alters filament dynamics also in this assay (relates to light scattering in Supplementary Figure S2C). Supplementary Figure S4B summarises statistics of the untagged AtubAB filament dynamics observed under the conditions used (Methods).

## DISCUSSION

Taken together (Figure 4D), our analyses showed that Heimdallarchaeia AtubAB tubulins are close relatives of eukaryotic tubulins and share with them many features and behaviours. They are encoded in the genome as a partly overlapping operon with a short intergenic region (Figures 1A & B, Supplementary Figures S1A - D). Expressed in *E. coli* and purified they form stable heterodimers, bind and hydrolyse GTP and polymerise in a crowding Polymerisation Buffer in a strictly GTP-dependent manner. Like tubulin, AtubAB polymerisation is also inhibited by the small molecule maytansine. When expressed in bacterial *E. coli* cells, AtubAB polymerise into 4-pf mini microtubules that are hollow and pseudo helical with a seam, both in cells and after purification, *in vitro*. Protofilaments alternate and are held together via the conserved M-loop. The monomers undergo a polymerisation-dependent cytomotive switch that enables polar dynamics with a fast plus and a slow minus end – supporting our previous supposition that this is a universal property of these polymers (Wagstaff et al., 2023). Finally, these polymers exhibit dynamic instability at their plus ends. Amazingly, all these properties are shared between AtubAB mini microtubules and eukaryotic microtubules, except their protofilament numbers: 4 vs 13.

What does this mean for the evolutionary relationship of AtubAB with microtubules? Let us first examine scenarios in which tubulin(s) forming microtubules were inherited vertically by early eukaryotes from their Asgard precursors. The presence of tubulin homologues in Asgard archaea is broad but patchy: tubulins are only found in approximately 1% of all surveyed Asgard archaeal genomes to date, in certain members of Lokiarchaeia, Odinarchaeia and Heimdallarchaeia (Hodarchaeales and Kariarchaeaceae). High rates of gene loss are common across all domains of life, and the scenarios in which tubulin(s) forming (mini) microtubules were inherited by early eukaryotes from their Asgard archaeal precursors may therefore imply losses of multiple paralogues in most Asgard archaeal lineages, as well as in most lokiarchaeia and heimdallarchaeia. These scenarios would therefore indicate complex combinations of duplications and losses across the Asgard archaea. Such scenarios could be supported by the co-existence of six stable paralogs in eukaryotes, and by the observed elevated rates of gene duplications in Asgard archaea compared to other archaea (Eme et al., 2023), which may suggest that Asgard archaeal gene histories could in some cases be as complex as those observed in early eukaryotic evolution (Vosseberg et al., 2021). Given the patchy distribution of these genes in Heimdallarchaeia, it is likely that there have been frequent horizontal transfer events of a subset of tubulin genes. However, the presence of tubulin homologues in a hodarchaeon and a kariarchaeon suggests the possibility that a distinct set of tubulin paralogues was present in the heimdallarchaeial ancestor of eukaryotes (Eme et al., 2023).

An alternative, second scenario consistent with the observed data, is the horizontal transfer of tubulin genes between Asgard archaeal lineages and early eukaryotic ancestors. However, there is no clear evidence of such events thus far. Currently, the lack of traces of synteny between Asgard archaeal lineages beyond the AtubAB operon or mobile elements in its vicinity prevents the identification of signals of joint transfer.

Exploring additional Asgard archaeal genome diversity will be key to identifying AtubABC orthologs or additional paralogous genes that will help reconcile the evolution of this gene family with the Asgard archaeal species tree. In the meantime, given current eukaryogenesis theory, we suggest that microtubules evolved in Asgard archaea before the formation of eukaryotes.

Because AtubA and AtubB are both more closely related to α tubulin, this suggests the intriguing hypothesis that the use of a similar AB heterodimer with a GTPase defective interface between the A and B subunits arose by a process of convergent evolution. This suggests that there is something special about using a heterodimer as building block of (mini and eukaryotic) microtubules.

Our findings raise important future questions: what is the function of mini microtubules in Asgard archaea? Our discovery of maytansine as a polymerisation inhibitor of AtubAB and recent advances with the cell biology of Asgard archaeal organisms (Radler et al., 2025) will help answering this question. Do molecular motor proteins exist in any prokaryote? This is an old question that so far has never received a positive answer. The discovery of (almost) non-twisting and stiff mini microtubules that display tubulin’s C-terminal α helices (helices H11/H12, Figure 3H) on the outside makes it more likely than ever that archaeal motor proteins may exist and they may, or may not be, related in sequence to eukaryotic motor proteins. Their existence would also need AtubAB mini microtubule dynamics to be regulated through stabilising (mini) microtubule-associated proteins (MAPs), whose possible existence will also need to be probed in archaea.

## DATA AVAILABILITY

The in-cell cryo-ET AtubAB mini microtubule map has been deposited in the Electron Microscopy Data Bank (EMDB) with accession number EMD-56154. The AtubAB cryo-EM structure has been deposited in the Protein Data Bank (PDB) under accession numbers PDB 9TQK and EMD-56153. The AtubAB cryo-EM heterodimer SPA structure has been deposited in the Protein Data Bank under accession numbers PDB 9TQJ and EMD-56152.

## ACKNOWLEDGEMENTS

We thank all members of the MRC Laboratory of Molecular Biology (LMB) Electron Microscopy Facility for help and support with data collection and Scientific Computing at the LMB for their support. We thank Tomos Morgan (LMB MS Facility) for performing intact mass spectrometry and Stephen McLaughlin (LMB Biophysics Facility) for performing the SEC-MALS analysis. We would like to thank the following LMB colleagues for help with initial experiments on other archaeal tubulins: Radu Aricescu, Veronica Chang, Fusinita van den Ent and Tim Nierhaus (all LMB). We would like to thank Buzz Baum (LMB) for helpful discussions. We would like to thank Mick Adriaansens (Wageningen University) and Mike Puijk (Utrecht University) for assistance with genomic analyses, and Jan Kees van Amerongen (Utrecht University) for his support and management of computational infrastructure.

This work was funded by the Medical Research Council, UK (U105184326 to JL, MC_UP_1201/31 to TAMB and MC_UP_1201/13 to ED), the Wellcome Trust (203276/C/16/Z, 227876/Z/23/Z and 227452/Z/23/Z to JL; 225317/Z/22/Z to TAMB), the Leverhulme Trust (Philip Leverhulme Prize to TAMB), the European Research Council (Consolidator and Advanced Grants 817834 and 101142180, respectively, to TJGE), the Dutch Research Council (VI.C.192.016 to TJGE and VI.Vidi.243.140 to DT), the Volkswagen Foundation (Life grant 96725 to TJGE), the Simons Foundation as part of the Moore-Simons Project on the Origin of the Eukaryotic Cell (grant 73592LPI to TJGE and BJB), the Spanish Ministerio de Ciencia e Innovación (CNS2023-145079 and PID2021-123399OB-I00 to MAO), the Australian Research Council (ARC Discovery Project DP230100769 to BPB) and The Trond Mohn Foundation through the Centre for Deep Sea research, University of Bergen to SLJ. This work made use of the Dutch national e-infrastructure with the support of the SURF Cooperative (EINF-2953 to TJGE).

## CONTRIBUTIONS

TJGE and JL conceived the study.

KEA, JED, FIM, S-JN, SLJ, BPB and BJB generated and analysed metagenomic data.

DT, JED, JV and SK performed sequence homology searches.

DT, JV and SK performed phylogenetic analyses.

JL expressed and purified AtubAB and did biochemistry.

JL and AvK performed structural biology, including cryo-EM. TAMB helped with cryo-EM screening.

MM performed cryo-ET and subtomogram averaging.

MAO and JL performed GTPase and maytansine experiments.

VPH and ED performed and analysed TIRF microscopy.

JL, TJGE and DT wrote the manuscript.

All authors edited the manuscript.

JL, BPB, BJB, ED, MAO, TAMB and TJGE provided supervision and funding.

## MATERIALS & METHODS

GTP lithium salt and GDP were purchased from Sigma, dissolved in water at 100 mM, pH adjusted with Tris (sat.) to pH 7 and aliquoted before freezing. GMPCPP was purchased from Jena Biosciences as a 10 mM solution and used as supplied. Maytansine was purchased from Merck (SML3451) and dissolved at 10 mM in DMSO (dimethyl sulfoxide) and kept frozen before use. The compounds paclitaxel (Alfa Aesar Chemicals), vinblastine, colchicine, podophyllotoxin and nocodazole (Sigma), epothilone A, plinabulin and pironetin (Med Chem Express), sabizabulin (TargetMol) were dissolved at 20 or 50 mM in DMSO and kept frozen before use. Gatorbulin was provided by Hendrick Luesch (University of Florida), discodermolide was provided by Ian Patterson (University of Cambridge), zampanolide was provided by Karl-Heinz Altmann (ETH Zurich), laulimalide was from Robert Keyzer and was purified from sponge biomass (Victoria University of Wellington, Ministry of Fisheries, Kingdom of Toga, permit F1/40/76/16 for collecBng the sponges). FcMaytansine was also provided by Karl-Heinz Altmann (ETH Zurich).

Two different versions of the general tubulin buffer BRB80 were used. For most experiments, BRB80 was 80 mM PIPES [piperazine-N,N’-bis(2-ethanesulfonic acid], 1 mM MgCl_2_, 1 mM EGTA [ethylene glycol-bis(2-aminoethylether)-N,N,N’,N’-tetraacetic acid], pH 6.9 [KOH]). 500 mM potassium glutamate were added to that, prior to final pH adjustment to 6.9, to produce the Polymerisation Buffer. TIRF experiments (below) used a slightly different version of BRB80, and this did not have scientific reasons.

### Identification of tubulin homologs and phylogenetic analysis

To recover a set of tubulin sequences for phylogenetic analyses, we started by performing broad sequence similarity searches using reference datasets, including COG5023, COG0206 and PF00091, against ENSEMBL and NCBI nr databases (versions September 2022). We removed redundancy using CD-Hit (-c 0.8) (Li et al., 2001), aligned these sequences using FAMSA (Deorowicz et al., 2016), trimmed using trimAl (-gt 0.003) (Capella-Gutiérrez et al., 2009) and reconstructed a phylogeny using FastTreeMP (Price et al., 2010). Using this tree, we selected sequences from an inclusive clade including all eukaryotic tubulin sequences. Additionally, we used the tubulin-family sequences from (Zaremba-Niedzwiedzka et al., 2017) to perform a sequence similarity search using PsiBLAST (-num_iterations 1 -evalue 0.001) (Altschul et al., 1997) against a local database including all public Asgard archaeal genomes (last updated: August 2023) and additional MAGs. We combined all these sequences in a single file, removed duplicates and all sequences shorter than 100 aa, aligned again using FAMSA, trimmed out gappy columns using trimAl (-gt 0.01), and reconstructed a phylogeny using IQ-TREE v2.2.2.6 (Minh et al., 2020) using LG+G4. We identified long branches using PhyKIT (lbs) (Steenwyk et al., 2021) and removed sequences with a score higher than 50. We then reconstructed one more tree by following the previous procedure: FAMSA, trimAl (-gt 0.01) and IQ-TREE under the model LG+G4. The resulting tree was used to manually classify eukaryotic sequences into the following groups: α tubulin, β tubulin, γ tubulin, δ tubulin, ε tubulin, and cryptic tubulins. We then down-sampled these sequences by running CD-Hit (Fu et al., 2012) (-c 0.55 for α, β and ε, -c 0.6 for γ, -c 0.7 for δ), to generate a balanced dataset). Separately, we recruited Asgard archaeal and verrucomicrobial (bacterial) BtubAB tubulins that clustered with eukaryotic tubulins and down sampled them separately using CD-Hit (-c 0.8), and sequences clustering with artubulins. To search for additional Asgard archaeal sequences published since our initial searches in 2023 (Appler et al., 2024; Langwig et al., 2025; Tamarit et al., 2024; Valentin-Alvarado et al., 2024), we combined the sequences in a single FASTA file, aligned it with MAFFT-linsi (Katoh et al., 2005), trimmed with trimAl (-gt 0.5) and used the resulting alignment to perform one more search against NCBI nr (July 2025) and a local database of Asgard archaeal genomes using PsiBLAST (-num_iterations 1 -evalue 1e-10).

To obtain a well-populated outgroup, we used CetZ sequences from our previous searches, plus a public sequence dataset by (Santana-Molina et al., 2023) and aligned them using FAMSA, together with the set of eukaryotic, Asgard archaeal and bacterial tubulins. We then trimmed this alignment using trimAl (-gt 0.5), removed gappy sequences (over 50% of gaps), and aligned with FastTreeMP (-lg). In the resulting tree, we selected sequences form a clade representing CetZ and removed redundancy using CD-Hit (-c 0.65). To obtain a sequence dataset for artubulins, we selected identified artubulins from (Santana-Molina et al., 2023) and from our initial searches above, aligned them using MAFFT-linsi and used this alignment for an additional sequence similarity search against NCBI’s NR database (February 2025) using PsiBLAST (-num_iterations -evalue 1e-100), and removed redundancy using CD-Hit (-c 0.8). To investigate the effect of combining different sets of sequences, we employed four datasets: one with all eukaryotic tubulins, Asgard archaeal tubulins, and bacterial tubulins; one adding artubulins; one adding CetZ; and one adding both artubulins and CetZ. For all these datasets, we aligned sequences using MAFFT-linsi, trimmed with trimAl (-gt 0.5 or -gt 0.1) and removed sequences with over 50% gaps. We evaluated model fit using ModelFinder (Kalyaanamoorthy et al., 2017) within IQ-TREE3 v3.0.1 (Wong et al., 2025) (command: iqtree3 -m MFP -mset WAG+C40, LG+C40, Q.pfam+C40, Q.pfam_gb+C60, WAG+C50, LG+C50, Q.pfam+C50, Q.pfam_gb+C60, WAG+C60, LG+C60, Q.pfam+C60, Q.pfam_gb+C60 -mrate G4 -mfreq ‘’ -n 0), and identified Q.pfam+C50+G4 as the best model according to the Bayesian Information Criterion in all instances. We then reconstructed phylogenies using IQ-TREE3 under this model, and an additional tree using the Posterior Mean Site Frequency (PMSF) approximation of this model (Wang et al., 2018), to reconstruct 100 standard bootstrap pseudo replicates. Some alignments were additionally trimmed during the reconstruction process by removing the columns with the 10% lowest log-likelihood values (“--robust-phy 0.9” in iqtree3) (Liu et al., 2025). Transfer Bootstrap Expectation (Lemoine et al., 2018) was calculated using RAxML-ng (Stamatakis, 2014). For visualisation, trees were rooted using madRoot (Tria et al., 2017) and plotted using figtree v1.4.4 (Rambaut, 2010). Maps to visualise the neighbourhoods around tubulin homologues were plotted using genoPlotR (Guy et al., 2010) including homology links identified with Diamond v2.1.9.163 (Buchfink et al., 2021) sequence similarity comparisons under default parameters.

### Cloning and expression of AtubAB in E. coli

Metagenomic sequencing revealed an operon in Kariarchaeaceae 81MaySF_Bin13 (Figures 1A & B, Supplementary Figure S1C) that on the translated protein sequence level showed significant sequence similarity to eukaryotic tubulins and bacterial BtubAB. We named these proteins Kari AtubA and AtubB and they were subsequently added by others to GenBank under accession numbers MDH5401500.1 and MDH5401501.1, respectively. AlphaFold 3 (Abramson et al., 2024) confirmed the tubulin fold of AtubA and B (Figure S1D), but also revealed that the amino acid sequence of AtubA as deposited in MDH5401500.1 is likely too short at the N-terminus since the first β strand is not completing the central β sheet in the N-terminal domain of the tubulin fold. Inspecting the nucleotide sequence revealed that there is likely a rare ATT start codon upstream and this produces the sequence MSEVVVV (replacing M1 in MDH5401500.1) that is related to other tubulins near the N-terminus as confirmed by multiple sequence alignments. In the modified operon, the ATT start codon also positions a likely RBS in a plausible distance to the first start codon (Supplementary Figure S1C). The resulting Kari *atubA* and *atubB* genes as used in this study are coloured blue and red in Supplementary Figure S1C. Expression of AtubAB in *E. coli* was facilitated by the construction of artificial bicistronic expression vectors. For this, the *atubA* and *atubB* genes were codon optimised for expression in *E. coli* (IDT) and a larger intergenic region with its own (non-overlapping) RBS was placed between them. Two constructs were made: AtubAB untagged and AtubAhBs, with AtubA His_8_-tagged and AtubB 2xStrep-tagged (each tag preceded by a TEV cleavage site). Overhangs for Gibson assembly into the expression vector pHis17 (for example Addgene plasmid #78201) were added to both constructs, resulting in the sequences listed in Supplementary Figures S2A and S2B, which were provided by IDT as gBLOCKS. Gibson assembly with a linearised version of pHis17 by PCR using the primers

> CGATCCGGCTGCTAACAAAGCCCGAAAGGA
>
> and CATATGTATATCTCCTTCTTAAAGTTAAAC

resulted in the two expression vectors, pHis17-AtubAB and pHis17-AtubAhBs, whose correctness was confirmed by Sanger sequencing. *E. coli* C41(DE3) cells were transformed with the two expression vectors and expression levels were tested in 10 mL 2xTY cultures containing 100 µM ampicillin at 37°C. After induction with 1 mM IPTG (isopropyl β-D-1-thiogalactopyranoside), cells were grown further at 37°C for 4-6 hours, harvested and taken up in 1 mL B-PER complete (Thermo Fisher Scientific) and incubated for 15 min at RT for lysis. Centrifugation at 20,000 xg at 4°C separated soluble lysates from insoluble pellets.

### Expression and Purification of His_8_- and Strep-tagged AtubAhBs

*E. coli* C41(DE3) cells were transformed with the pHis17-AtubAhBs expression vector and grown first in 200 mL and then 6L 2xTY supplemented with 100 µM ampicillin at 37°C / 200 rpm. After addition of 1 mM IPTG, cells were grown for another 5 hrs at 37°C / 200 rpm, harvested by centrifugation and stored at -70°C. 500 mL of Buffer A (50 mM Tris, 200 mM NaCl, pH 7.5) with some DNase I and six cOmplete protease inhibitor tablets (Roche) were added to the cells to resuspend. Cells were lysed by a single pass through a Constant Systems cell disruptor at 35 kPSI. Ultracentrifugation in a Beckman Ti45 rotor at 35,000 rpm for 1 h was used to remove insoluble material. The lysate was pumped through two HisTrap HP nickel columns (Cytiva) at 2 mL/min. Washing and elution was done at 7 mL/min using Buffer A and Buffer B (1 M imidazole [only], pH 7.0) and involved steps of 2%, 5%, 10%, 30% and 100% Buffer B. AtubAhBs eluted mostly in the 10% fractions, which were checked by SDS-PAGE and Coomassie staining. Protein was precipitated by adding 100% (sat.) ammonium sulphate to 50% (sat.) final concentration. The precipitate was isolated by centrifugation at 45,000 xg at 4°C for 20 min. The pellet was resuspended in 2 mL of Buffer A and subjected to size exclusion chromatography (SEC) using a Sephacryl S200 16/60 (Cytiva) column in Buffer A at 1 mL/min. Fractions were checked by SDS-PAGE and those containing AtubAhBs were pooled and frozen into aliquots at -70°C. Final concentration (without concentrating) was 6 g/L as determined with a UV spectrometer and a calculated extinction coefficient. Over 100 mg of protein were obtained. The identity of the proteins was confirmed by intact mass spectrometry after TEV cleavage overnight (0.35 g/L TEV protease): AtubAh: 49,035 Da (theoretical, no M1: 49,036 Da); AtubBs: 47,033 Da (theoretical, no M1: 47,034 Da). The un-cleaved AtubAhBs product with the His_8_- and Strep-tags still present was used in this study.

### Expression and purification of untagged AtubAB

*E. coli* C41(DE3) cells were transformed with the pHis17-AtubAB expression vector and grown first in 200 mL and then 6 L 2xTY supplemented with 100 µM ampicillin at 37°C / 200 rpm. After addition of 1 mM IPTG cells were grown for another 5 hrs at 37°C / 200 rpm, harvested by centrifugation and stored at -70°C. 500 mL of Buffer C (20 mM Tris, pH 8.0) with some DNase I and six cOmplete protease inhibitor tablets (Roche) were added to the cells to resuspend. Cells were lysed by a single pass through a Constant Systems cell disruptor at 35 kPSI. Ultracentrifugation in a Beckman Ti45 rotor at 35,000 rpm for 1 h was used to remove insoluble material. The lysate was pumped through a HiPrep Q XL 16/10 anion exchange column (Cytiva) at 10 mL/min. Proteins were eluted at the same speed with a gradient to 50% of Buffer D (Buffer C + 1 M NaCl) against Buffer C. Fractions were checked by SDS-PAGE and Coomassie staining. Protein was precipitated by adding 100% (sat.) ammonium sulphate to 50% (sat.) final concentration. The precipitate was isolated by centrifugation at 45,000 xg at 4°C for 20 min. The pellet was resuspended in 3 mL of Polymerisation Buffer (BRB80 with 500 mM potassium glutamate: 80 mM PIPES [piperazine-N,N’-bis(2-ethanesulfonic acid], 1 mM MgCl_2_, 1 mM EGTA [ethylene glycol-bis(2-aminoethylether)-N,N,N’,N’-tetraacetic acid], 500 mM potassium glutamate, pH 6.9 [KOH]) and subjected to size exclusion chromatography (SEC) using a Sephacryl S200 16/60 (Cytiva) column in Polymerisation Buffer at 1 mL/min. Fractions were checked by SDS-PAGE and those containing AtubAB were pooled, concentrated with a Vivaspin 20 10 kDa molecular weight cut off (MWCO) concentrator (Sartorius) to ∼30 g/L (as determined by UV with calculated extinction coefficient) and frozen into aliquots at -70°C. Over 100 mg of protein were obtained. The identity of the proteins was confirmed by intact mass spectrometry: AtubA: 48,097 Da (theoretical, no M1: 48,098 Da); AtubB: 46,095 Da (theoretical, no M1: 46,095 Da).

### Size-exclusion chromatography with multi-angle light scattering (SEC-MALS)

SEC coupled with multi-angle static light scattering (SEC-MALS) was performed using an Agilent 1200 series LC system with an online Dawn Helios ii system (Wyatt Technology) equipped with a QELS+ module (Wyatt Technology) and an Optilab rEX differential refractive index detector (Wyatt Technology). Untagged AtubAB at 3 g/L was injected onto a Sephadex S200 Increase HR10/300 (Cytiva) size exclusion column pre-equilibrated in Polymerisation Buffer (BRB80 + 500 mM potassium glutamate; 80 mM PIPES, 1 mM MgCl_2_, 1 mM EGTA, pH 6.9 [KOH]). The light scattering and protein concentration at each point across the peaks in the chromatograph were used to determine the absolute molecular mass from the intercept of the Debye plot using Zimm’s model as implemented in the ASTRA v7.3.2.19 software (Wyatt Technology). To determine inter-detector delay volumes, band-broadening constants and detector intensity normalisation constants for the instrument, BSA (bovine serum albumin) was used as a standard prior to sample measurements.

### GTPase assay

GTP hydrolysis by untagged AtubAB was monitored through the detection of the release of inorganic phosphate with a malachite green assay, following a previously published procedure (Kodama et al., 1986). Samples contained 25 µM protein (dimer, 2.35 g/L) in Polymerisation Buffer (BRB80 with 500 mM potassium glutamate, pH 6.9 [KOH]) at 25°C.

### Critical concentration determination

10 g/L (94 µM dimer) of untagged AtubAB in BRB80 with 500 mM potassium glutamate Polymerisation Buffer were diluted 3 times 1.5-fold in Polymerisation Buffer in 200 µL volumes. 100 µL samples were pre-spun 20 min at 50,000 rpm for 20 min at 20°C in a TLA100 rotor (Beckman). 2 mM GTP was added, and the samples were incubated for 30 min at RT. Samples were then centrifuged at 50,000 rpm for 20 min at 20°C in the same TLA100 rotor (Beckman). The supernatants were withdrawn; pellets were washed with 50 µL Polymerisation Buffer (no nucleotide) and were resuspended in 100 µL of gel loading buffer over 1 h. Samples from supernatants and pellets were prepared in gel-loading buffer (1:10 dilution) for SDS-PAGE and subsequent Coomassie staining. Band intensities were integrated with BioRad’s Image Lab software and plotted.

### Pelleting assays

Figure S2E: 2.3 g/L (25 µM dimer) of AtubAB in BRB80 with 500 mM potassium glutamate Polymerisation Buffer were assembled in the presence of 2 mM GTP with 27.5 µM of LAU (laulimalide), PTX (paclitaxel), ZAM (zampanolide), DIS (discodermolide), EPO (epothilone), MAY (maytansine), PIR (pironetin), GAT (gatorbulin), VIN (vinblastine), COL (colchicine), POD (podophyllotoxin), SAB (sabizabulin), NOC (nocodazole) and PLI (plinabulin) or the DMSO (dimethyl sulfoxide) control. After 2 h at 37°C, samples were centrifuged at 60,000 rpm for 30 min at 37°C in a TLA100 rotor (Beckman). The supernatants were withdrawn, and pellets were resuspended in same volume of Polymerisation Buffer. Samples from supernatants and pellets were prepared in gel-loading buffer for SDS-PAGE. Figure 2G: as above but 6 g/L (63 µM dimer) pre-spun (20 min at 50,000 rpm for 20 min at 20°C in a TLA100 rotor [Beckman]) AtubAB were mixed with 50 µM MAY (maytansine) and incubated at RT for 30 min, before spinning at 50,000 rpm for 20 min at 20°C in a TLA100 rotor (Beckman). The DMSO and compound reactions contained 1 mM GTP. Figure 2E: as Figure 2G but 10 g/L (94 µM dimer) pre-spun (20 min at 50,000 rpm for 20 min at 20°C in a TLA100 rotor [Beckman]) AtubAB were mixed with 2 mM GDP or GTP, or 0.5 mM GMPCPP (Jena Bioscience) and incubated at RT for 30 min before spinning at 50,000 rpm for 20 min at 20°C in a TLA100 rotor (Beckman).

### Nucleotide content determination

A solution containing 25 µM (2.4 g/L) of untagged AtubAB in BRB80 with 500 mM potassium glutamate Polymerisation Buffer was used. The sample was sedimented (100,000 xg for 40 min at 25°C) and nucleotide from supernatant and pellets were extracted with cold 0.5 N HClO_4_ (with guanosine as internal standard). After 10 min at 4°C denatured protein was removed by centrifugation (12,000 xg for 10 min at 4°C). Aliquots were neutralised by the addition of 1/6 volume 1 M K_2_HPO_4_, 0.5 M acetic acid and 1/6 volume of 3 M KOH. For HPLC analysis of the nucleotide content, precipitated KClO_4_ was removed by centrifugation (12,000 xg for 10 min at 4°C) and nucleotides were separated by isocratic reverse-phase ion pair HPLC (Supelcosil LC-18-DB) in buffer 0.2 M K_2_HPO_4_, 4 mM tetrabutylammonium, 0.1 M acetic acid, pH 6.7 as described: (Seckler et al., 1990). Two independent experiments were run and produced similar results.

### Maytansine binding affinity determination

Fluorescence anisotropy competition experiments and analyses were performed as previously described (Díaz and Buey, 2007), with minor modifications. The binding constant of the fluorescent FcMaytansine (maytansine with fluorescein attached) probe to purified untagged AtubAB was determined by fluorescence anisotropy titration at 25°C in black Nunc 96-well, flat-bottom microplates and in a final volume of 200 μL. 90 nM FcMaytansine in assay buffer (15 mM PIPES-KOH, pH 7.0, 1 mM EGTA supplemented with 1.5 mM MgCl_2_ and 0.1 mM GTP) was titrated with increasing amounts of AtubAB up to 1 µM. The anisotropy and fluorescence values of the free (r = 0.035 ± 0.001) and bound (r = 0.256 ± 0.005) states were determined in the absence and presence of 1 µM AtubAB, respectively. The anisotropy was measured using an iD5 microplate reader (Molecular Devices). Samples were equilibrated at 25°C prior to measurement. Excitation and emission wavelengths were set to 485 and 435 nm, respectively. The read height and the gain were previously adjusted using both fluorescein and FcMaytansine. To calculate the binding constant of FcMaytansine, the fractional saturation, νb, of the probe had to be determined. Binding of FcMaytansine to AtubAB was measured through changes in its anisotropy. The anisotropy of a mixture of free and bound FcMaytansine can be expressed as

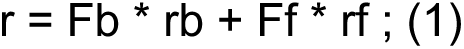

where r is the measured anisotropy, Ff and Fb are the fractional fluorescence intensities of free and bound FcMaytansine, respectively. rf is the anisotropy of the free FcMaytansine, and rb is the anisotropy of the bound FcMaytansine. Because there is a 2.6 ± 0.2-fold difference between the fluorescence intensity of the bound and free FcMaytansine, the fluorescence fraction is weighted by the change in fluorescence intensity according to

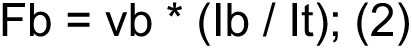

where Ib is the fluorescence intensity of bound FcMaytansine and It is the total fluorescence intensity. Given these two equations

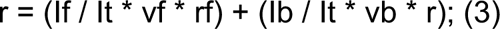

where If is the fluorescence intensity of free FcMaytansine and νf is the fractional saturation of free FcMaytansine. Since the sum of the fractions of free (νf) and bound (νb) ligand is 1 and

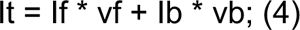

Employing equations 1, 3 and 4, we obtain the fractional saturation of FcMaytansine needed to calculate the binding constant of its binding to the site and of a ligand by competition with a probe of known binding constant.

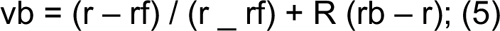

with R representing the ratio between the fluorescence intensity of the bound and free species (R = Ib/If).

Once the fractional saturation of FcMaytansine in the experiments is known, the free concentration of AtubAB and FcMaytansine in the titration assay can be calculated from the difference between the bound and the total concentration of the probe. The free concentration of AtubAB was calculated from the difference in the total and bound concentration of FcMaytansine assuming a stoichiometry of 1:1. The values of fractional saturation of FcMaytansine bound vs. the free AtubAB concentration were used to determine a binding constant at 25°C using SigmaPlot v13 (Systat Software).

Then, samples containing 90 nM FcMaytansine and 90 nM untagged AtubAB in assay buffer were titrated with increasing amounts of a competitor at 25°C in black Nunc 96-well, flat-bottom microplates and in a final volume of 200 μL. Fluorescence anisotropy measurements were carried out as described above. The binding constant of maytansine K(m) can be determined from the known values of the binding constant of FcMaytansine K(Fc) and the total concentrations of binding sites, by solving the equations:

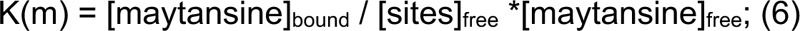

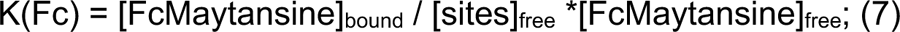

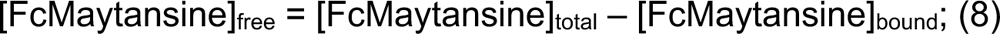

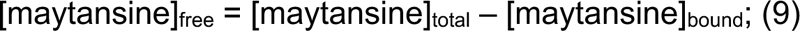

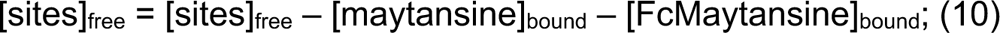

The system was solved using EQUIGRA5 (Díaz and Buey, 2007), using the binding constant of FcMaytansine and a stoichiometry of 1:1 for the AtubAB-FcMaytansine complex

### 90° light scattering assays

90° light scattering was measured on a Cary Eclipse fluorescence spectrometer (Agilent Technologies). A non-stirred cuvette was used: 10 mm 28F/MS-Q10 (Spectrology), taking 900 µL of sample. Settings were as follows: 360 nm excitation and emission wavelengths, 23°C reaction temperature, 5 nm slits, 1 s averaging, photomultiplier setting (PMT) low and 700 min total collection time. 6 g/L (64 µM dimer) of tagged AtubAhBs or untagged AtubAB in BRB80 with 500 mM potassium glutamate, pH6.9 (KOH) Polymerisation Buffer were used in 900 µL volumes. 44 µM of nucleotides were used to start the reactions. MAY (maytansine) at 10 mM in DMSO was added to 200 µM final concentration.

### Focussed ion beam (FIB)-milling for lamellae production

To produce samples of sufficiently thin ice for subtomogram averaging (STA), grids of *E. coli* C41(DE3) cells over-expressing untagged AtubAB (plasmid pHis17-AtubAB) were prepared for cryo-FIB milling. Cells were grown in 2xTY media to OD_600_ ∼1 before induction with 1 mM IPTG for several hours until > OD_600_ 3 was reached. Cells were pelleted at 1,800 xg for 5 min, before resuspending in fresh media to ∼ OD_600_ 60 (viscous). Quantifoil Au R2/2 200 mesh grids were glow discharged using a PELCO easiGlow at 25 mA for 45 s. Samples were plunge frozen using a Vitrobot Mk IV (Thermo Fisher Scientific) at 21°C with humidity control turned off. 8 µL of concentrated *E. coli* sample were applied to the front faces of the grids, which were then back blotted for 6 s (blot force 10) with the aid of a custom-made Teflon ring in the shape of a Vitrobot blot paper. Plunge-frozen grids were then clipped and loaded onto a Thermo Fisher Scientific Aquilos 2 Dual Beam cryo-FIB/SEM for automated milling and polishing. Lamellae positions were assigned using MAPS software (Thermo Fisher Scientific), while milling was performed using AutoTEM software (Thermo Fisher Scientific) with our own optimised milling procedure. Grids were coated with a layer of organo-platinum using a Gas Injection System (GIS) for 40 s, followed by 30 s of sputtering with inorganic platinum to reduce charging effects during milling. A milling angle of 12° was chosen for all sites. Milling of lamellae using the gallium beam (all at 30 kV) was fully automated, aside from initiation of the final polishing step and was carried out in several steps of incrementally reduced milling currents and lamella widths. Rough milling was carried out at 1 nA (rectangular pattern) with a starting lamella width of 10 µm, followed by successive steps at 0.3 nA, 0.1 nA and 50 pA. Each site was milled ‘site wise’ (as opposed to ‘step wise’) overnight, ensuring up to 30 lamellae were milled up to the final polishing step in the morning. A final polish at 30 pA was performed the following morning for the best 20 lamellae, ensuring samples remained as freshly polished as possible, before transferring out of the Aquilos 2 cryo-FIB/SEM. Final lamella thicknesses were targeted to 110 nm, with a final width of 6 μm. See Figure S3A for actual lamellae thicknesses as determined by subsequent cryotomography.

### Electron cryotomography

Lamellae prepared by cryo-FIB milling were transferred to a Thermo Fisher Scientific Krios G4 300 kV Transmission Electron Microscope (TEM), equipped with a Falcon 4i detector, a SelectrisX energy filter and a cold field emission gun. Data were collected using Tomography 5 software (Thermo Fisher Scientific), with search maps of lamellae used to target regions for collection. A magnification of 83kx (1.514 Å/pixel) was used to collect tilt series bidirectionally. A starting stage tilt angle of 12° followed by 3° tilt increments to -42° (for the first branch), before completing the second branch from 12° to 66°. The samples were exposed for 1.27 s across 8 frames per tilt image, with a dose accumulation of 4.1 e^-^/Å^2^/tilt or 151.7 e^-^/Å^2^ across all 37 tilts. An objective aperture of 100 µm and an energy filter slit width of 10 eV was selected and centred immediately prior to data collection. A target defocus range of -1.5 to -3.5 µm was applied across the data collection.

### Subtomogram averaging (STA) of AtubAB mini microtubules in E. coli

Tilt series data was imported into the Relion 5 STA pipeline (Burt et al., 2024). Tilt series were motion corrected using Relion’s implementation of MotionCorr2 (Zheng et al., 2017) with 5 x 5 patches, followed by CTF estimation with CTFFIND-4.1 (Rohou and Grigorieff, 2015). Bad tilts were manually removed using the Napari-based viewer (Ahlers et al., 2023; Gaifas et al., 2023) in Relion 5, before tilt series alignment in Relion 5 with AreTomo2 (Zheng et al., 2022) using an estimated tomogram thickness of 120 nm and correcting for tilt angle offset (because of the FIB milling angle). Tomograms were reconstructed with unbinned pixel dimensions of 4,000 x 4,000 x 2,000 at 10 Å/pixel and denoised using CryoCARE (Buchholz et al., 2019). For all tomograms, the sample thicknesses were measured to be at or below 200 nm using the software Geollama (Rosalind Franklin Institute, 2025). Filament picking was carried out on denoised tomograms using the Relion 5 Napari picking viewer allowing annotation of filament paths. Particle poses were determined using ‘relion_python_tomo_get_particle_poses’ with unknown filament polarity. All subsequent processing steps used Relion 5 2D tilt stacks for particles. Initial poses were extracted at bin 3 (4.542 Å/pixel), with a maximal dose accumulation of 60 e^-^/Å^2^ and a box size of 70 binned pixels (318 Å). Due to the uncertainty around the filament composition (number of protofilaments), an initial model-free helical refinement against a cylinder without imposed symmetry was performed, revealing a 4-stranded filament. With the subunit spacing along the helical axis being approximately 40 Å (the canonical longitudinal tubulin repeat), a helical rise of 10 Å was estimated to allow for 4 subunits per 40 Å. With the direction of the twist unknown, two refinements were conducted in tandem against a cylinder, one with a 90° twist and another with a -90° twist. The negative twist refinement presented with a better final map and a substantially better angular accuracy. Thus, a final ‘initial’ refinement was performed on the extracted coordinates with a -90° twist and 10 Å rise using the negative twist map as a reference (low-pass filtered to 30 Å). A subsequent masked refinement with symmetry searching was performed, yielding a map with an estimated resolution of 9 Å (Nyquist limit at bin 3) and helical parameters of -89.86° twist and 10.29 Å rise. These particles were then re-extracted at bin 2 with a box size of 106 pixels (321 Å) and a maximum dose accumulation of 60 e^-^/Å^2^. Another masked refinement was performed following removal of duplicate particles resulting in an 8.22 Å resolution map (gold standard FSC cutoff 0.143). 3D classification was performed to remove suboptimal particles, leaving 16k particles for another round of 3D refinement, this time with Blush regularisation (Kimanius et al., 2024). This resulted in a map with an estimated resolution of 7.5 Å. These particles were then extracted at bin 1 for a round of Relion 5 Tomo refinement cycles consisting of CTF refinements and Bayesian polishing. Following an improvement in resolution from each step, a new bin 1 volume was reconstructed to benefit from the improvements gained. Polishing was performed with per-particle motion switched on and per-frame 2D deformation estimations. These steps improved the resolution of reconstructed volumes to 6.4 Å (FSC 0.143 criterion). Further refinements and classifications did not improve resolution much more and so the final map chosen was that reconstructed from the last round of Bayesian polishing. Local resolution was estimated in Relion 5 and the final reconstruction was sharpened using EMReady (He et al., 2023). See Supplementary Figure S3D for an overview of STA processing.

### Negative staining electron microscopy

Untagged AtubAB in Polymerisation Buffer (BRB80 with 500 mM potassium glutamate, pH 6.9 [KOH]) was diluted to 6.6 g/L with Polymerisation Buffer and 3 mM GTP (or 3 mM GDP, or 0.6 mM GMPCPP) were added. After 1 h incubation at RT, the sample was rapidly diluted to 0.3 g/L with Polymerisation Buffer and 5 µL were applied to 50 s glow-discharged, carbon-coated EM grids (5-6 nm amorphous carbon film, 400 mesh copper, CF400-CU, Electron Microscopy Sciences) and incubated for 30 s. After blotting off the protein solution with filter paper, 5 µL of 2 % (w/v) uranyl acetate solution were applied three times and blotted off immediately with filter paper each time. Images were taken at various magnifications on a Phillips Tecnai T12 Spirit, operated at 120 kV and equipped with a Gatan Orius CCD detector.

### Cryo-EM (SPA) sample preparation

For AtubAB filaments, untagged AtubAB in Polymerisation Buffer (BRB80 with 500 mM potassium glutamate, pH 6.9 [KOH]) was diluted to 7 g/L with Polymerisation Buffer and 4 mM GTP were added. After incubating the sample at RT for 1 h, the sample was further diluted to 0.25 g/L with Polymerisation Buffer and used immediately. For vitrification, a Leica EM GP2 automatic plunger was used. Temperature for plunging was 18°C, humidity was 99% (rel.) and a blot time of 5 s was used. 3.5 µL of sample were added to 60s glow-discharged Quantifoil Au R2/2 200 mesh grids and blotted, before vitrification in liquid ethane kept at -180°C. For the un-polymerised AtubAB heterodimer, untagged AtubAB in Polymerisation Buffer was diluted to 0.5 g/L in 50 mM HEPES (4-(2-hydroxyethyl)-1-piperazineethanesulfonic acid), 100 mM KCl, 5 mM Mg acetate, 1 mM EGTA, pH 7.7 [KOH]. For vitrification, a Vitrobot Mark IV (Thermo Fisher Scientific) automatic plunger was used. Temperature for plunging was 8°C, humidity was 100% (rel.) and a blot time of 3.5 s at blot force 10 was used. 3.5 µL of sample were added to 60s glow-discharged Quantifoil Au R1.2/1.32 300 mesh grids and blotted, before vitrification in liquid ethane kept at -180°C.

### Cryo-EM (helical SPA) structure determination of AtubAB mini microtubules

All processing of micrographs was performed in Relion 5 (Scheres, 2012), unless otherwise stated. 4,349 movies were collected at 1.222 Å/pixel using EPU software (Thermo Fisher Scientific) on a Krios G4 microscope equipped with a Falcon 4i camera, a SelectrisX energy filter and a cold field emission gun (Thermo Fisher Scientific), operated at 300 kV. The movies were combined and motion corrected using Relion’s own implementation before contrast transfer function (CTF) estimation using CTFFIND4 (Rohou and Grigorieff, 2015). Because single filaments were rare (because of bundling), manual picking was used to produce 61k particles, extracted 42 Å along each filament in boxes of 266 pixels. Initial helical parameters were determined by using CryoSPARC 4 (Punjani et al., 2017) helical refinement without providing helical parameters or a model. This produced a clear 4-stranded density without difficulty, with the helical parameters twist = -89° and rise = 10.5 Å. These parameters describe an arrangement that must be pseudo-helical for tubulin-like heterodimers because the rise per near-full turn (-356°) is 42 Å, which is the same distance from one subunit in an alternating AtubAB (…ABABAB…) protofilament to the next (A to B) and not to the same type, two subunits away (A to A). This means, the structure must have a seam where A contacts B and *vice versa* (see Figure 2G for a schematic of this). In other words, helical processing with the parameters determined will average A and B subunits onto each other. Obtaining such an averaged map was done first using Relion 5 helical Refine3D with the helical parameters. Subsequent Class3D, CTFRefine and Polish jobs, followed by another round of helical Refine3D produced a fully symmetrical (A and B not distinguished) density at 3.1 Å resolution (twist = -89.84°, rise = 10.46 Å; FSC criterion 0.143). To resolve the A and B subunits in the map, resolving the 8-fold redundancy (4-fold around the tube and 2-fold along the tube) was tried by first symmetry expanding 8-fold with Relion 5’s relion_particle_symmetry_expand command, with the helical parameters, and subsequent Class3D without alignment into different numbers of classes and different T values - this failed. Instead, Class3D without alignment, with a mask around a single dimer succeeded, as indicated by the finding that the one major difference in structure between AtubA and B that AlphaFold 3 predicted (residues 102-111) was resolved almost completely (Figure 2J). This did not resolve the seam position but makes it possible to demonstrate that the structure solved is made of AtubA and AtubB in alternating protofilaments. The symmetry dictates that such a structure must have a seam. Because the Class3D job is not gold standard, phenix.auto_sharpen (Liebschner et al., 2019) was used to produce a final density for the heterodimer in the AtubAB filament. Relion’s ModelAngelo (Jamali et al., 2024) was used to build atomic models into the map, which went well, completely and automatically building AtubA on top and AtubB at the bottom of the dimer, without human intervention. The model was checked and refined further with rounds of refinement with phenix.real_space_refine and manual building in MAIN (Turk, 2013). Phenix.model_vs_map reported a final model to map resolution of 3.1 Å (FSC criterion 0.5). To obtain a complete atomic model, the AtubAB mini microtubule, 4 refined dimers were placed in the symmetrised map and symmetry expanded with the symmetry twist = 1.31° (8 * -89.84° + 720°), rise = 83.68 Å (8 x 10.46 Å). For an overview of the helical cryo-EM processing please consult Figure S3D.

### Cryo-EM (SPA) structure determination of un-polymerised AtubAB

All processing of micrographs was performed in Relion 5 (Scheres, 2012). 10,328 movies were collected at 0.955 Å/pixel using EPU software (Thermo Fisher Scientific) on a Krios G4 microscope equipped with a Falcon 4i camera, a SelectrisX energy filter and a cold field emission gun (Thermo Fisher Scientific), operated at 300 kV. The movies were combined and motion corrected using Relion 5’s own implementation before contrast transfer function (CTF) estimation using CTFFIND4 (Rohou and Grigorieff, 2015). Particles were picked using Topaz (Bepler et al., 2019) and its general model produced ∼1 million particles which were extracted in boxes of 210 pixels. Multiple rounds of Class2D reduced particle numbers to 380k, revealing clear side and top views of the AtubAB heterodimers (Figure S3H). Refine3D with a heterodimer reference model (AtubAB filament structure as determined above) filtered to 20 Å resolution produced a good map that revealed secondary structure elements but also highlighted that parts of AtubA were disordered. Rounds of Class2D and Class3D without alignment reduced particle numbers to 190k, at which point CTFRefine and Polish, followed by Refine3D produced an even better map. One round of Class3D without alignment found 125k good particles that after a last Refine3D and Postprocess produced the final heterodimer density at 3.5 Å resolution (FSC criterion 0.143). The previously obtained AtubAB heterodimer atomic structure was placed in the density and refined and adjusted with rounds of phenix.real_space_refine (Liebschner et al., 2019) and manual building in MAIN (Turk, 2013). For an overview of the processing please consult Figure S3I.

### AtubAB labelling for TIRF microscopy

AtubAB filament labelling with fluorescent dyes was performed by adopting protocols used for bacterial BtubAB previously (Deng et al., 2017). Aliquots of AtubAB were thawed and diluted to 40 μM (dimer, 3.8 g/L) in Polymerisation Buffer (for TIRF, only: 80 mM K-PIPES, 500 mM potassium glutamate, 1 mM EGTA, 5 mM MgCl_2_, 2 mM GTP, pH 6.9 [KOH]) and incubated at RT for 40 min to allow for filament polymerisation. The reaction was centrifuged at 100,000 x*g* in a TLA 120.2 rotor (Beckman) for 10 min at 20°C, the supernatant was discarded and the pellet resuspended in Polymerisation Buffer, adjusting the AtubAB concentration to 40 μM. For fluorescence labelling, the filaments were then used to resuspend a full tube of either Alexa Fluor™ 647 NHS Ester (Invitrogen A20006) or Atto 488 NHS ester (Sigma-Aldrich 41698-1MG-F). For biotin labelling, ∼2 mg of EZ-Link™ Sulfo NHS-LC-LC-Biotin (Thermo Fisher Scientific 21343) were resuspended in Polymerisation Buffer and added to the filaments at a final concentration of 400 μM (10-fold molar excess). The reactions were incubated at RT for 30 min with gentle rocking, then quenched by adding 1 mM Tris-HCl, pH 7.0 and incubating for 5 min at RT. After incubation, the samples were transferred into polycarbonate ultracentrifuge tubes (Beckman), pre-filled with 150 µL of Cushion Buffer (Polymerisation Buffer supplemented with 60% [v/v] glycerol). Cushion Buffer was used to ensure the removal of the non-reacted free dye. The tubes were then centrifuged at 100,000 x*g* in a TLA 120.2 rotor (Beckman) for 10 min at 20°C. The supernatant was discarded, and the pellet was resuspended (depolymerised) in Glutamate-free Buffer (80 mM K-PIPES, 1 mM EGTA, pH 6.9 [KOH]. The filaments were incubated at RT for 1 h to ensure their depolymerisation. Subsequently, the reaction was centrifuged again at 100,000 x*g* in a TLA 120.2 rotor (Beckman) for 20 min at 20°C. The supernatant was recovered; the protein concentration and the labelling efficiency were estimated using a nanodrop spectrometer (Thermo Fisher Scientific) and calculated extinction coefficients. The proteins were mixed to make two different stocks: AtubAB seed stocks were prepared at 50 μM final concentration (20% fluorescent-AtubAB, 10% biotinylated-AtubAB) in Polymerisation Buffer supplemented with 5 mM GMPCPP (Jena Bioscience NU-405S); free (un-polymerised) AtubAB-dimers were prepared at a final concentration of 50 μM (20% fluorescent-AtubAB) in Glutamate-free Buffer. Both solutions were aliquoted in 2 μL, immediately flash-frozen after mixing and stored for future experiments. Tagged AtubAhBs was added unlabelled.

### AtubAB TIRF Microscopy

22 x 22 mm glass coverslips (NEXTERION, Schott) were incubated for at least 48 hrs at RT with gentle agitation in a 1:10 (w/w) mix of mPEG Silane (30 kDa, PSB-2014, Creative PEGWorks) and PEG-Silane-Biotin (3.4 kDa, Laysan Bio) at a final concentration of 1 g/L in 96 % (v/v) ethanol and 0.2 % (v/v) HCl. On the day of each experiment, the coverslips were washed with ethanol and ultrapure water, dried with a nitrogen gas gun, and assembled into an array of flow cells on mPEG-Silane passivated slides using double-sided tape (Adhesives Research AR-90880, precisely cut with a Graphtec CE6000 cutting plotter). The chamber was first perfused with BRB80 buffer (TIRF, only: 80 mM K-PIPES, 1 mM EGTA, 5 mM MgCl_2_, pH 6.9 [KOH]), then NeutrAvidin (25 µg/mL, Thermo Fisher Scientific) was added to the chamber and incubated for 5 min, then washed out with 10 chamber volumes of BRB80. In the meantime, an aliquot of BtubAB-seeds was thawed, 2 μL of Polymerisation Buffer (BRB80 + 500 mM potassium glutamate, pH 6.9 [KOH]) was added and the reaction was incubated at room temperature for 5 min to allow for filament polymerisation. The reaction was then diluted to 30 μL in Imaging Buffer (80 mM K-PIPES, 200 mM potassium glutamate, 1 mM EGTA, 5 mM MgCl_2_, 2 mM GTP, 0.1 g/L potassium casein, 0.1 g/L BSA, 40 µM DTT, 64 mM D-glucose, 160 µg/mL glucose oxidase, 20 µg/mL catalase, pH 6.9 [KOH]) and added to the chambers. After 5 min, the chamber was quickly washed with 50 μL of imaging buffer and dynamic filaments were elongated from the seeds by injecting into the chamber a solution of 20% labelled AtubAB at a final concentration of 4 μM, supplemented with 5 mM GTP. Experiments with AtubAhBs were performed following the same procedure, but the final concentrations in the chamber was 5 μM, composed of 20% fluorescent-AtubAB, 60% unlabelled AtubAB and 20% AtubAhBs. The microscope stage was kept at RT. Images were collected every 5-10 s in a custom spinning disk/total internal reflection fluorescence (TIRF) microscope previously described in (Planelles-Herrero et al., 2025).

## Supplement

### SUPPLEMENTARY FIGURES

**Figure S1A.**
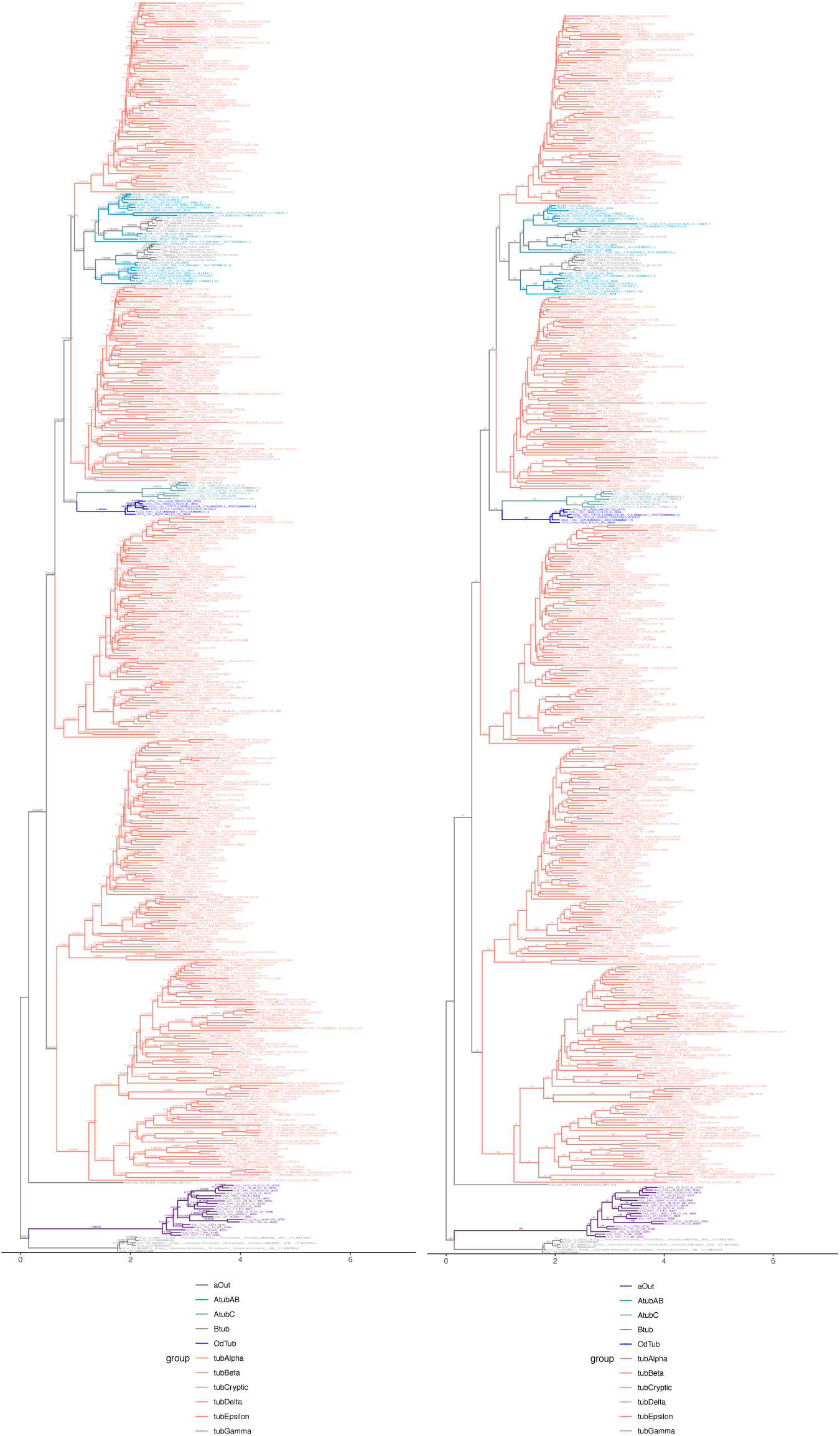
Full phylogeny corresponding to Figure 1A. The full tree is shown twice, with Transfer Bootstrap Expectation scores as branch support values shown on the left and Felsenstein Bootstrap Proportions on the right.

**Figure S1B.**
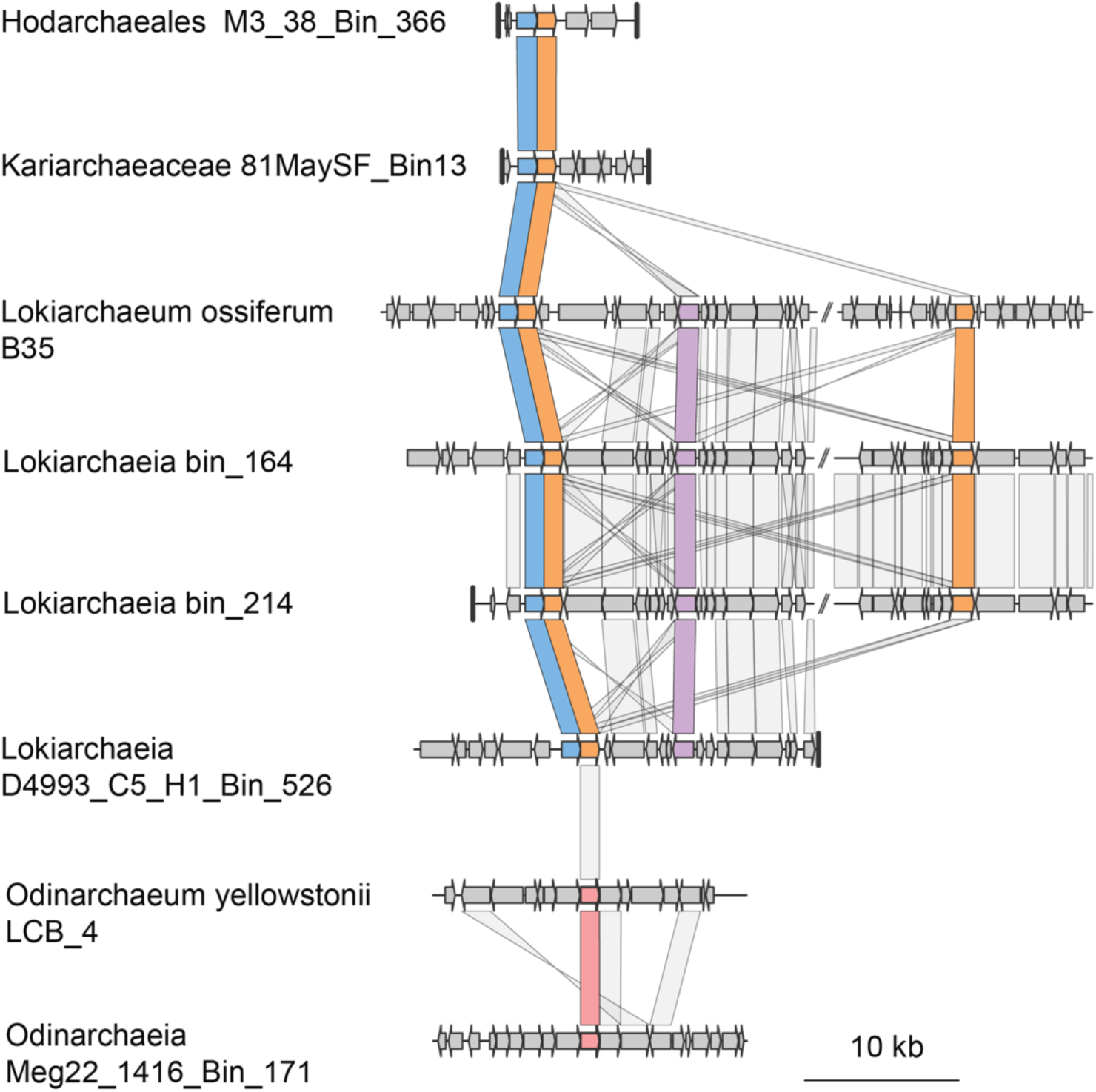
Gene maps of Asgard archaeal tubulins and their genomic regions(7 kb up and downstream). Genes are represented with arrows, and dark vertical lines indicate contig boundaries. Comparison lines indicate homology as identified by Diamond sequence similarity searches (Buchfink et al., 2021).

**Figure S1C.**
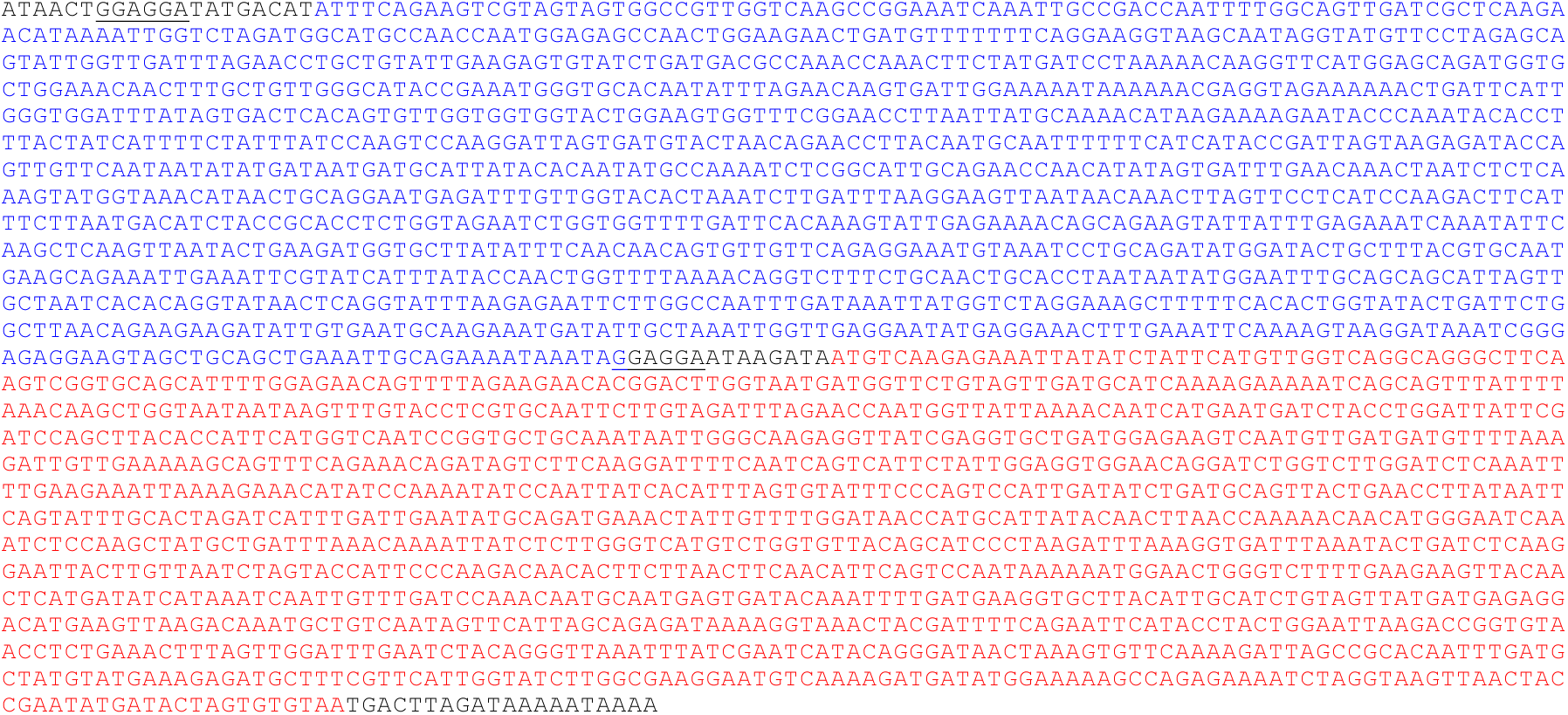
Kariarchaeaceae 81MaySF_Bin13 AtubAB operon metagenomic sequence (as shown schematically in Figure 1B). Note that *atubA* was extended upstream towards the rare ATT start codon because the deposited predicted protein sequence (GenBank: MDH5401500.1) is too short for the tubulin fold (as determined with AlphaFold 3), where the residues missing from the deposited sequence are needed to form the first β strand of the protein. The start codon was previously misassigned, most likely. Blue: *atubA*, red: *atubB.* Putative ribosome binding sites (RBSs) have been underlined.

**Figure S1D.**
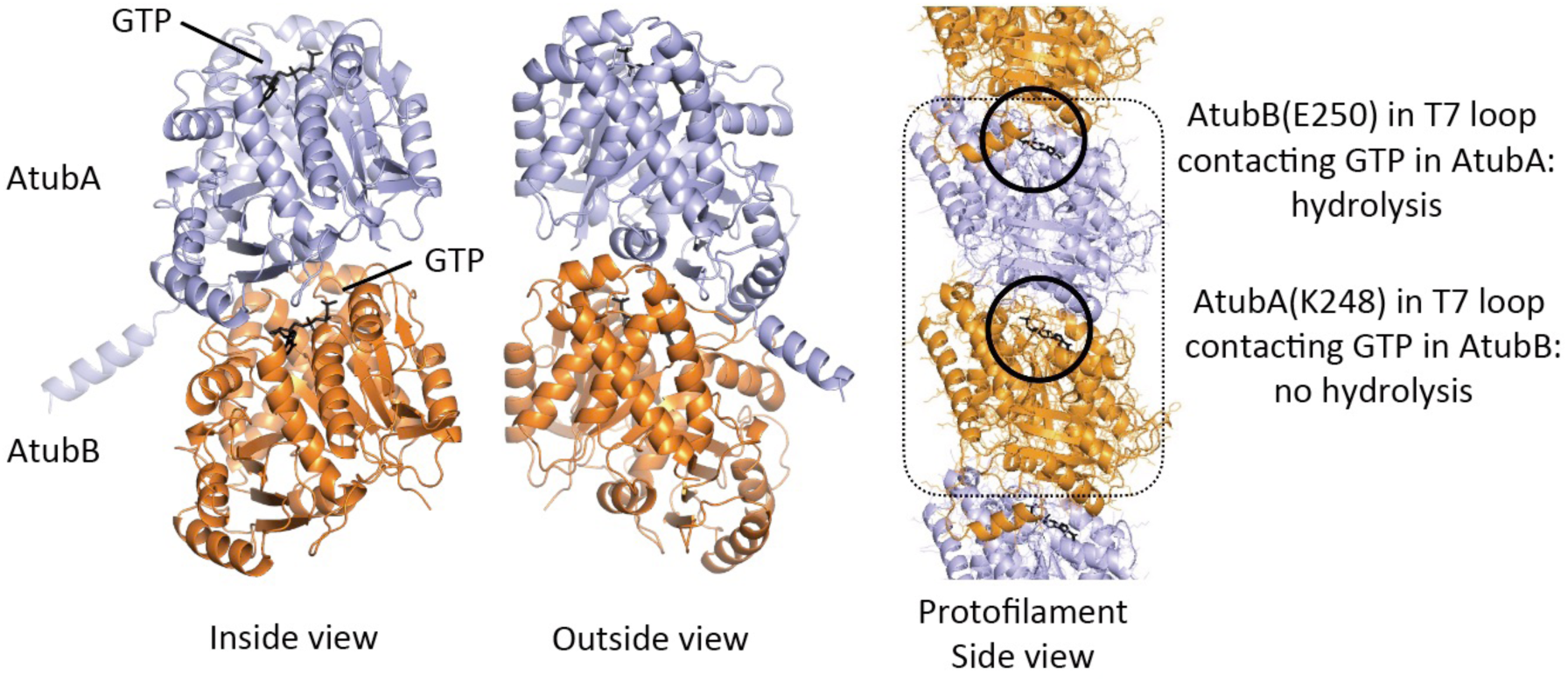
AlphaFold 3 prediction of AtubAB and stable heterodimer formation. AlphaFold 3 predicts AtubAB to be of the tubulin fold and to form heterodimers, and alternating ABAB protofilaments. Inspecting the AtubA and AtubB T7 loops in such a protofilament reveals that likely only one GTPase site is active, the one in AtubA (blue).

**Figure S2A.**
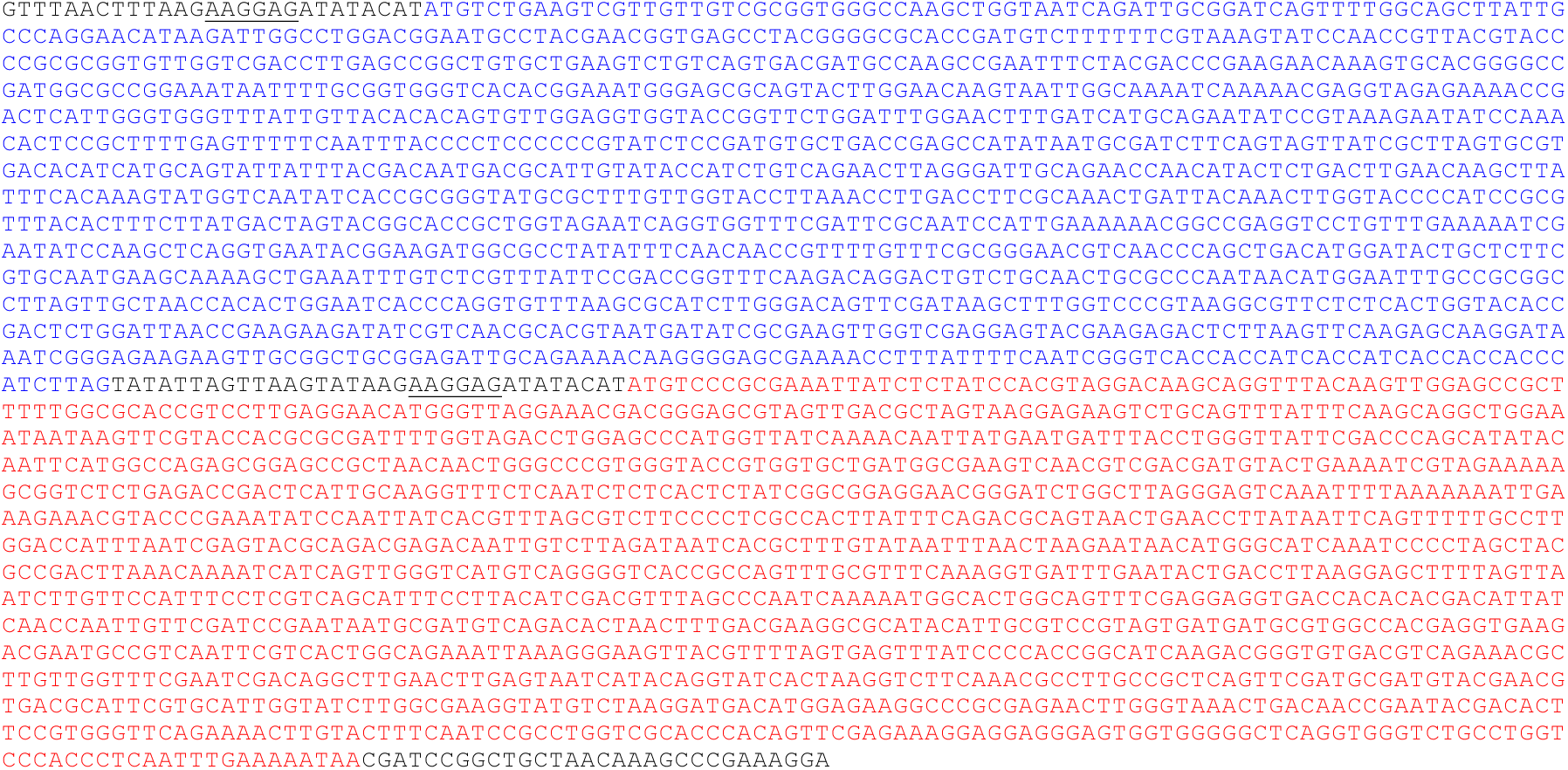
AtubAB His_8_- and Strep-tagged expression construct (AtubAhBs), codon optimised for expression in *E. coli*. Blue: *atubAh*, red: *atubBs.* Sequence of the gBLOCK used to make expression plasmid pHis17-AtubAhBs. RBSs have been underlined.

**Figure S2B.**
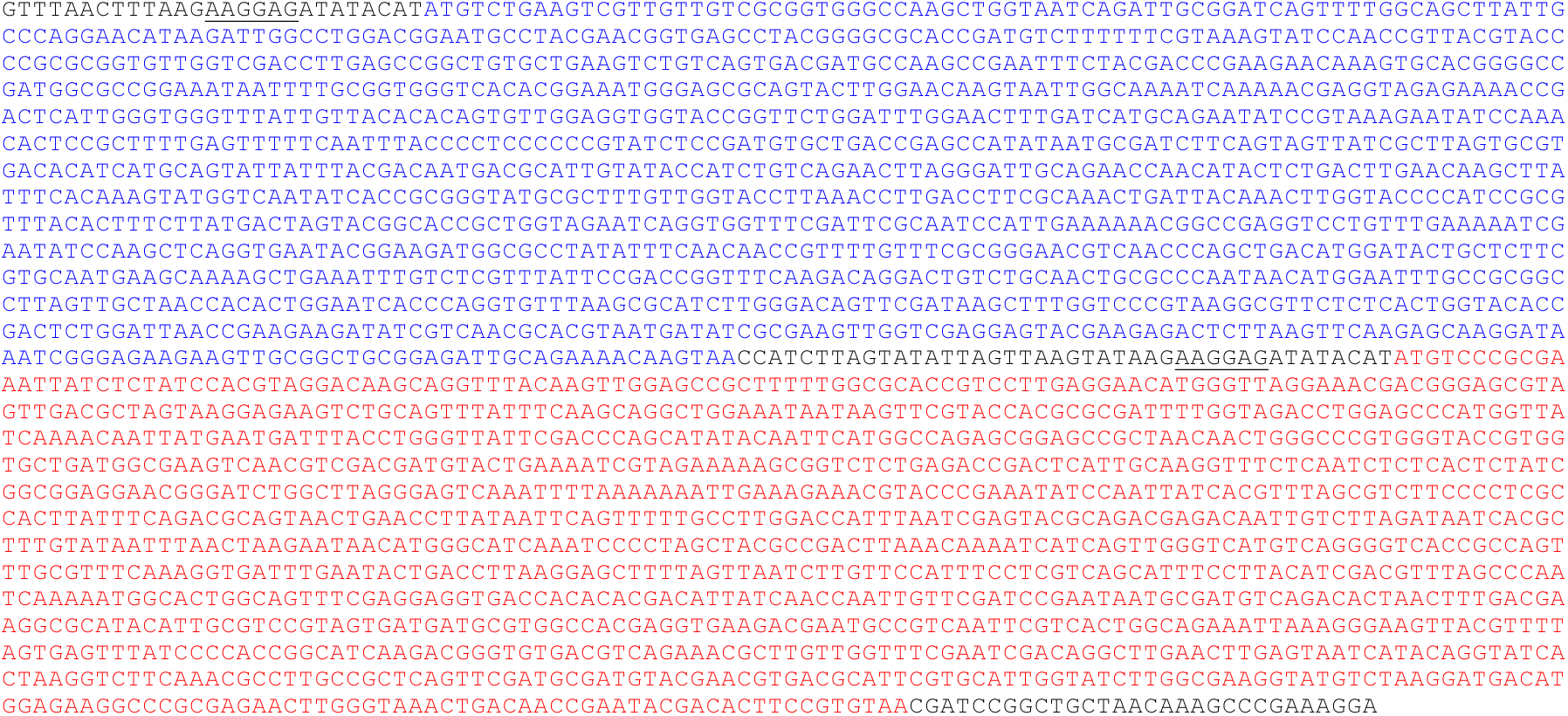
AtubAB untagged expression construct, codon optimised for expression in *E. coli*. Blue: untagged *atubA*, red: untagged *atubB.* Sequence of the gBLOCK used to make expression plasmid pHis17-AtubAB. RBSs have been underlined.

**Figure S2C.**
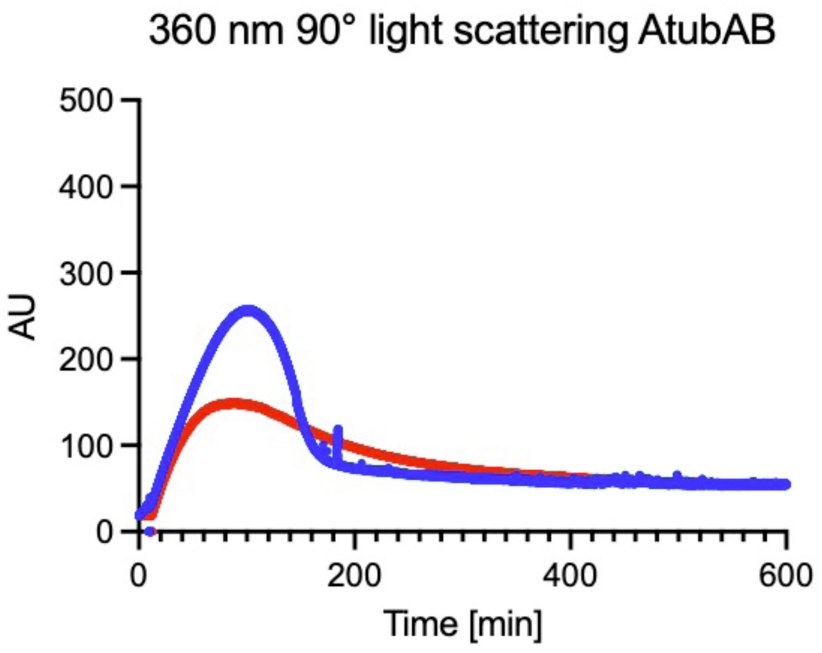
Tagged AtubAhBs 360 nm 90° light scattering. Experiment the same as in Figure 2F, but using tagged AtubAhBs protein instead (blue trace), which appears in this assay to polymerises more and depolymerises faster than untagged AtubAB (red trace).

**Figure S2D.**
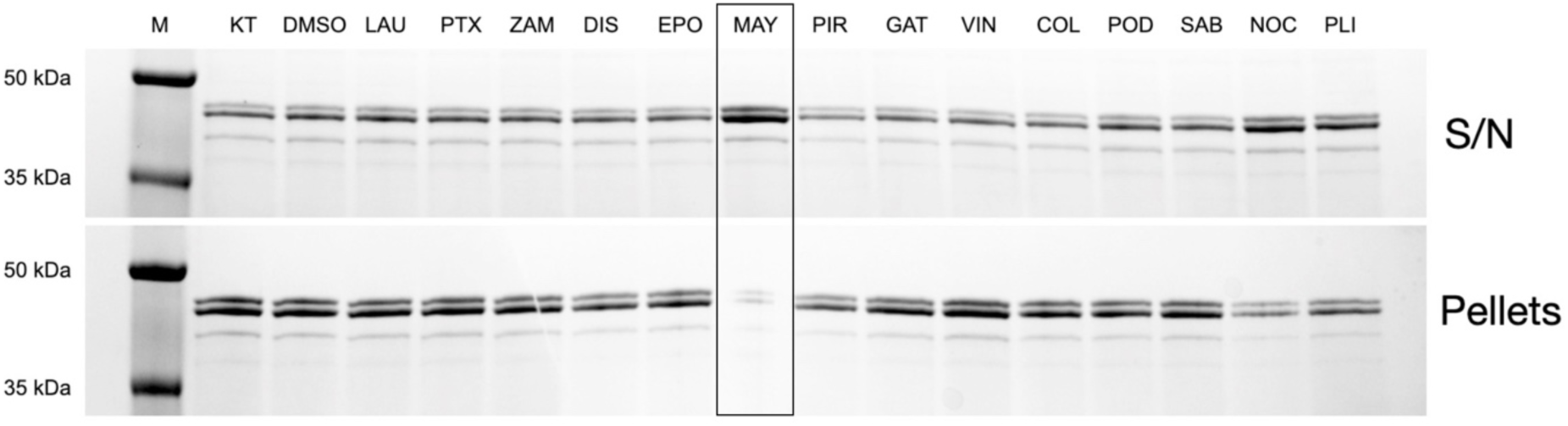
AtubAB pelleting assays with tubulin-directed small-molecule compounds. A panel of tubulin-directed drugs and inhibitors were tested against AtubAB polymerisation in a pelleting assay. Maytansine (MAY), a tubulin polymerisation inhibitor effectively inhibits pelleting of AtubAB in this assay. The same effect is shown in different experiments in Figure 2G and 2H. Key: KT (Kari AtubAB, nothing-added control), DMSO (dimethyl sulfoxide control), LAU (laulimalide), PTX (paclitaxel), ZAM (zampanolide), DIS (discodermolide), EPO (epothilone), MAY (maytansine), PIR (pironetin), GAT (gatorbulin), VIN (vinblastine), COL (colchicine), POD (podophyllotoxin), SAB (sabizabulin), NOC (nocodazole) and PLI (plinabulin). S/N, supernatant.

**Figure S2E.**
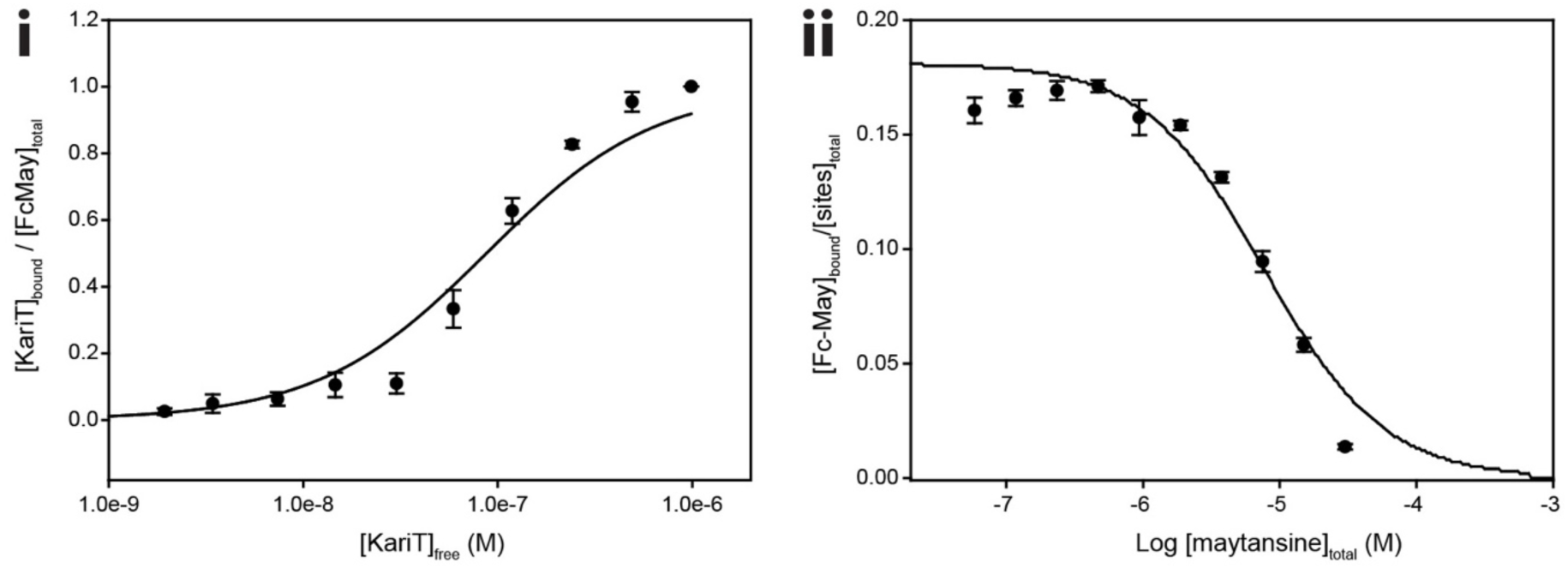
Affinity of maytansine for AtubAB. **i)** Binding affinity measurement using the fluorescently labelled maytansine compound FcMaytansine (Methods). The affinity was determined to be K_d_= 87 ± 17 nM (mean ± SEM, *n* = 4, see Methods). ii) Competition experiment to determine the AtubAB affinity of unlabelled maytansine. K_d_ = 5.1 ± 0.2 µM.

**Figure S2F.**
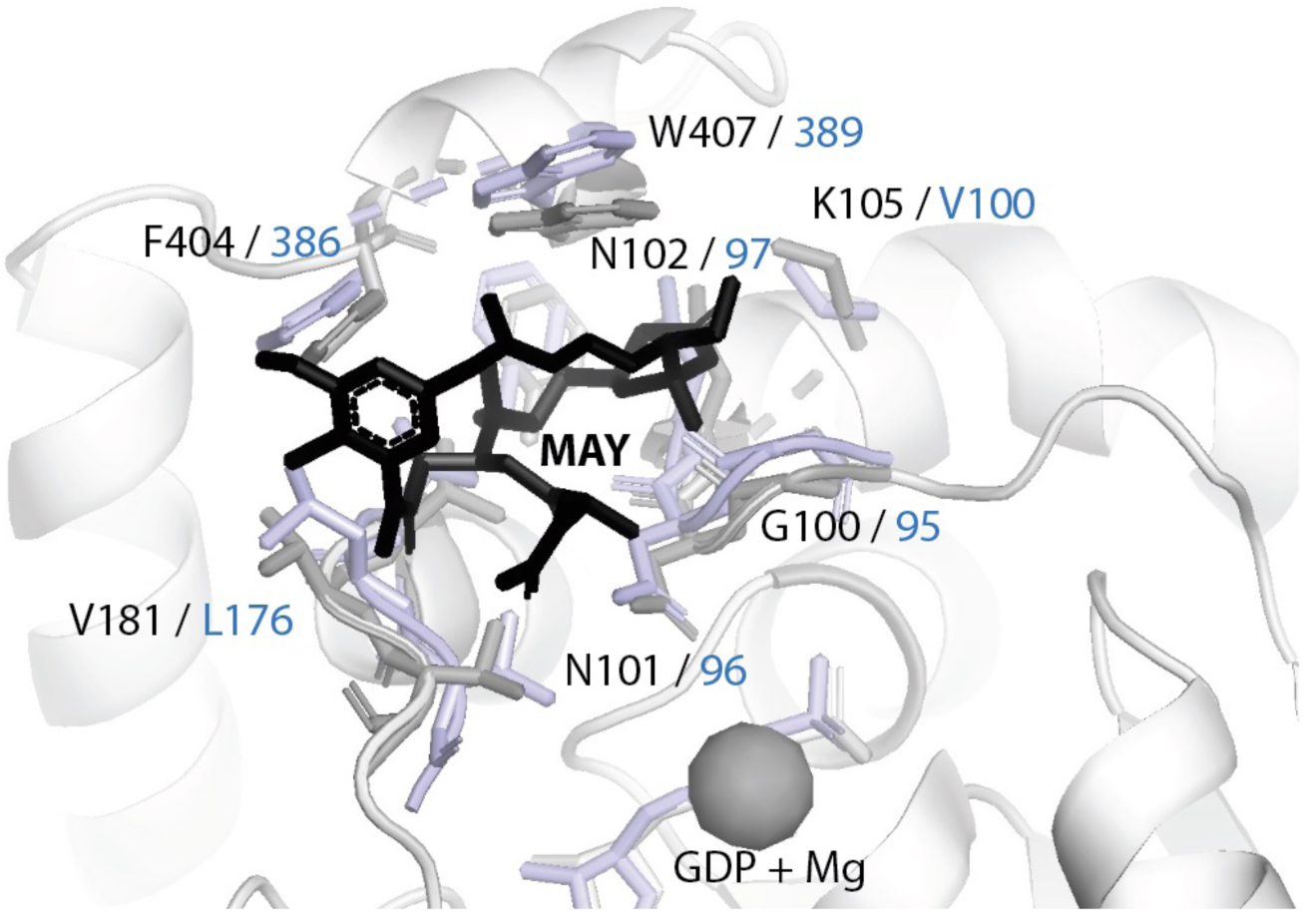
Superposition of tubulin complexed with maytansine and AtubAB AlphaFold 3 model. β tubulin:maytansine complex (PDB 4TV8) (Prota et al., 2014) superimposed on AtubA. The binding pocket is highly conserved - all interacting residues are identical except β tubulin(K105)/AtubA(V100) and β tubulin(V181)/AtubA(L176).

**Figure S3A.**
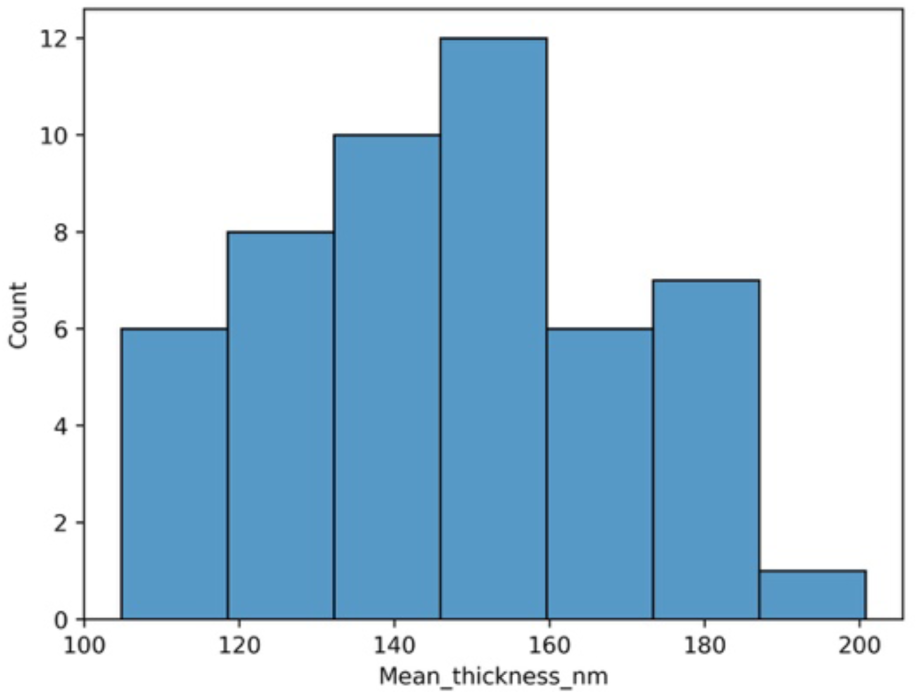
Distribution of lamellae thicknesses after focussed ion beam (FIB)-milling of *E. coli* C41(DE3) cells over expressing untagged AtubAB. The lamellae were used to perform electron cryotomography with subsequent subtomogram averaging to solve the *in situ* structure of AtubAB (Figures 3B, S3D). Thicknesses were determined with the software GeoLlama after tomography (Rosalind Franklin Institute, 2025). Note that 50 lamellae were prepared but three were discarded before reconstructing them (47 in total, Supplementary Table 1).

**Figure S3B.**
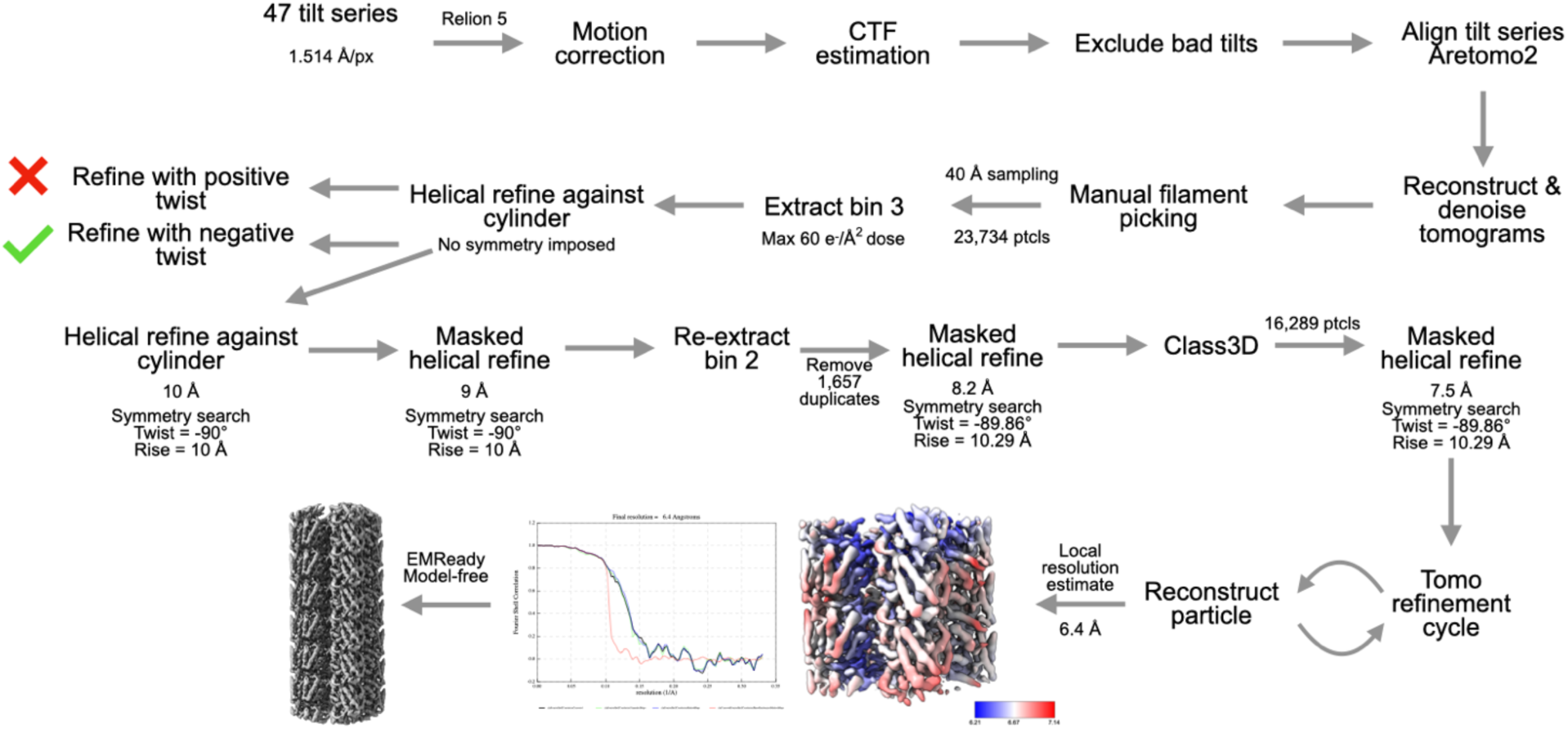
Cryo-ET / subtomogram averaging (STA) processing scheme. *E. coli* C41(DE3) cells over expressing untagged AtubAB and FIB-milled.

**Figure S3C.**
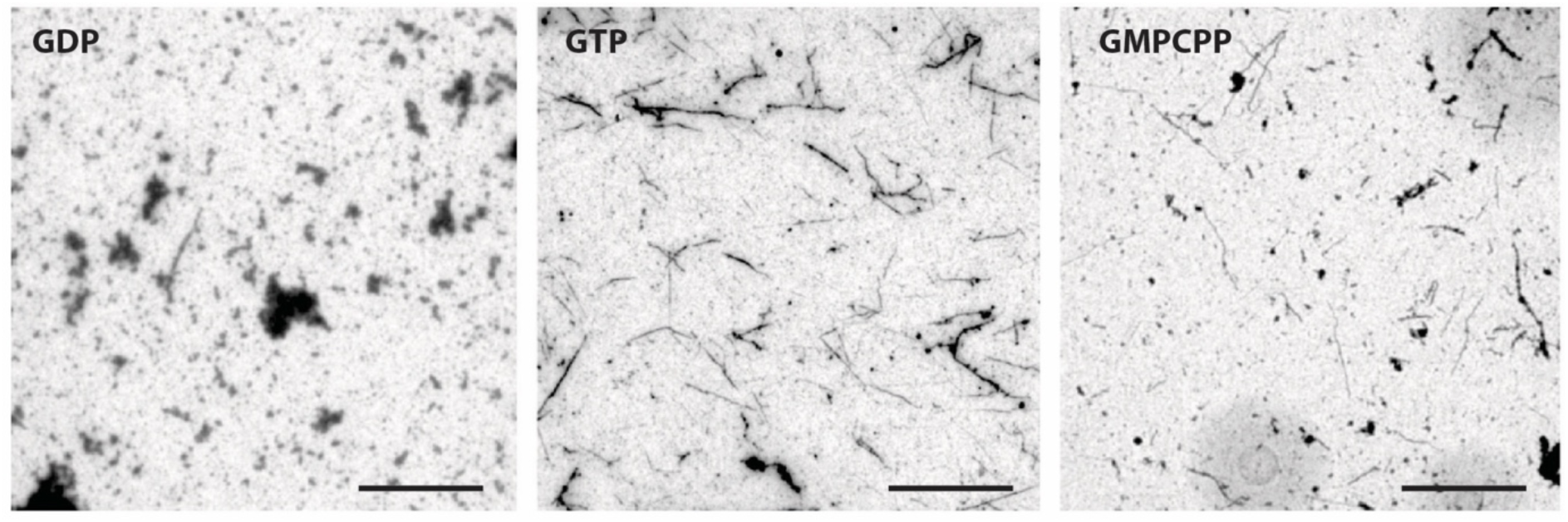
Low-magnification negative staining of AtubAB. Polymerisation performed in BRB80 + 500 mM potassium glutamate Polymerisation Buffer with GDP, GTP and GMPCPP nucleotides (scale bars = 5 µm). GTP and GMPCPP-containing reactions show many filaments. See also the corresponding pelleting experiment in Figure 2E.

**Figure S3D.**
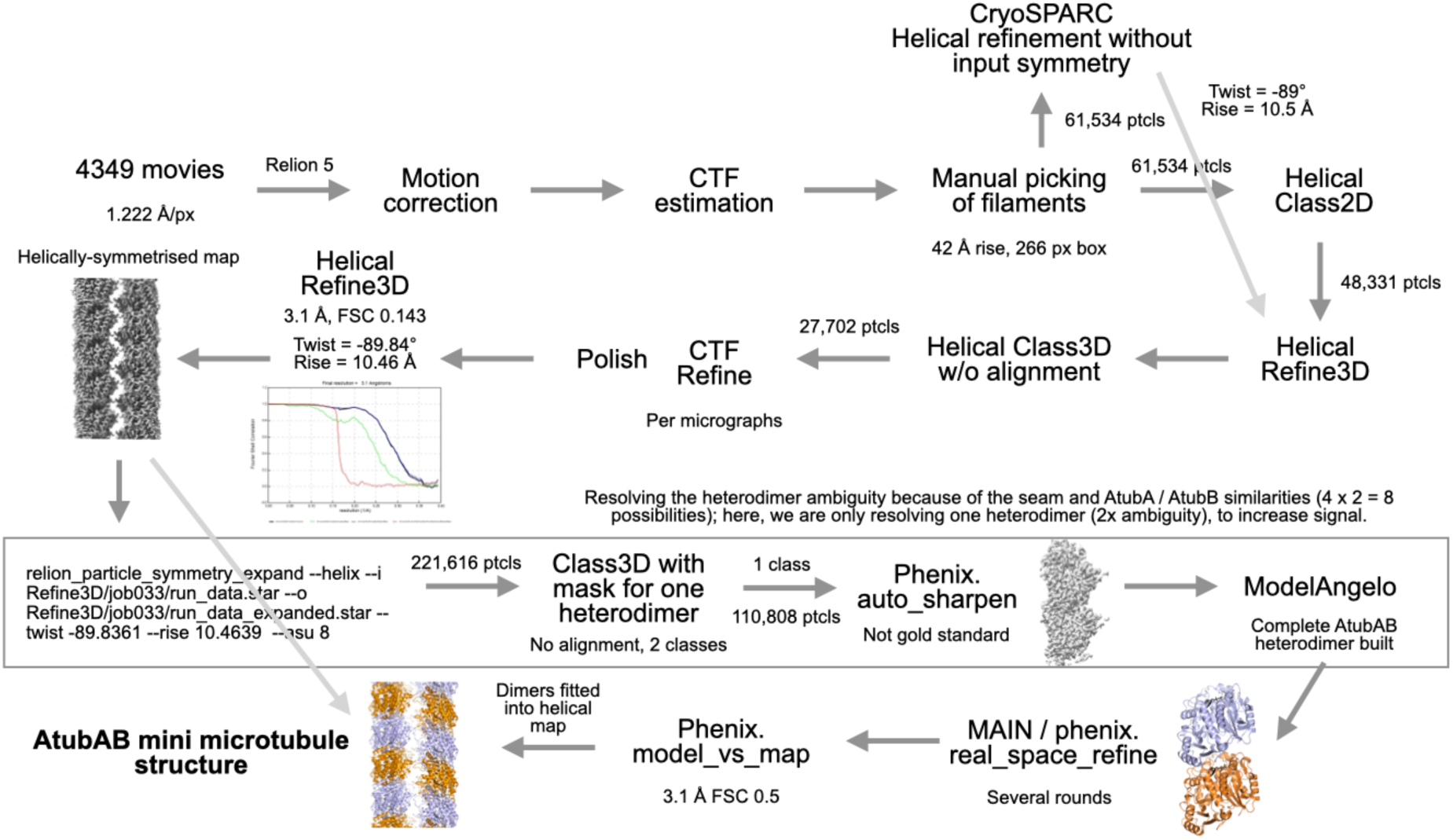
Cryo-EM processing scheme: helical processing and symmetry breaking of AtubAB mini microtubules.

**Figure S3E.**
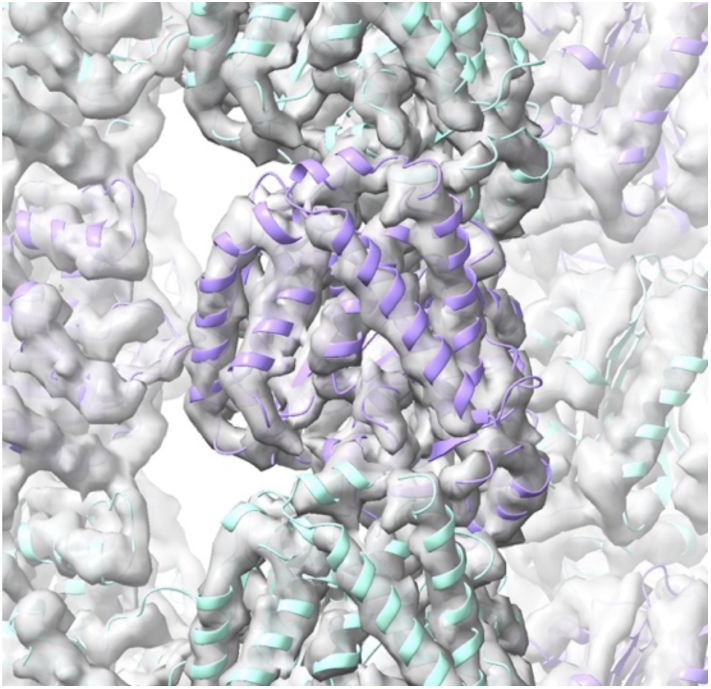
AtubAB model-free and subtomogram-averaged *in situ* map (Figure 3B), fitted with the *in vitro* cryo-EM atomic model (Figure 3F). Note that the colours are different here because A and B could not be identified in the 6.4 Å resolution map. Helices are resolved; sheets are not, in line with the resolution estimate.

**Figure S3F.**
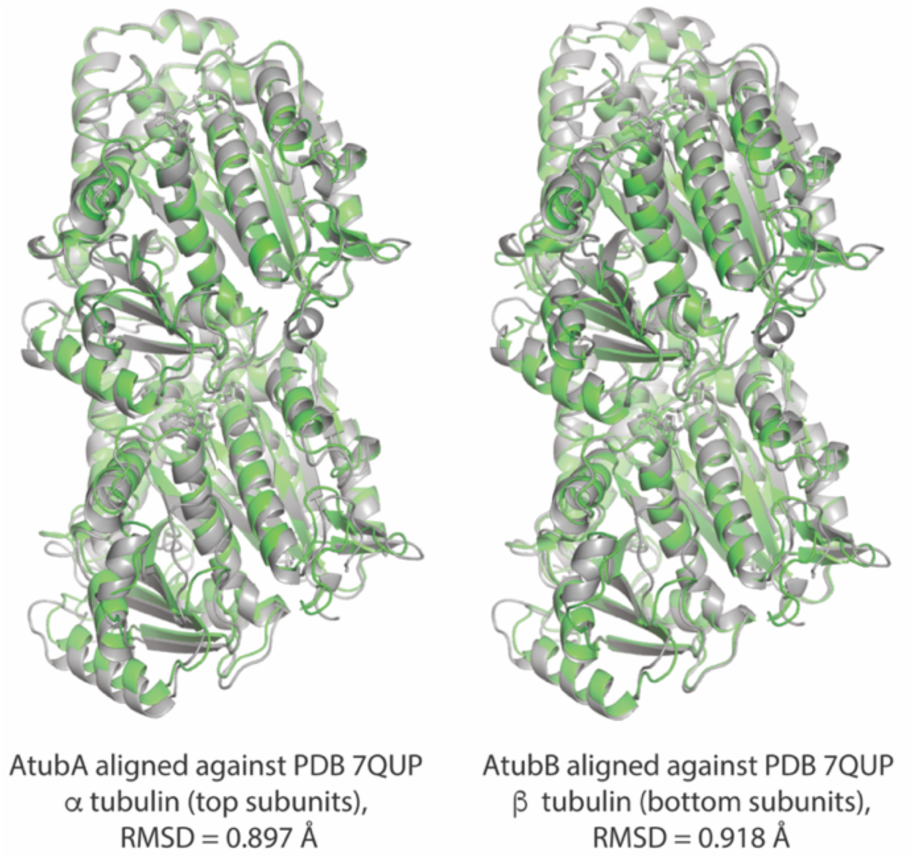
Superposition of AtubAB mini microtubule heterodimer, as determined in this study, with eukaryotic αβ tubulin in a microtubule. Left: AtubA (top) superimposed on PDB 7QUP (*Drosophila melanogaster* microtubule) β tubulin (2251 out of 2650 residues superimposed in PyMOL, RMSD = 0.897 Å) (Wagstaff et al., 2023); right: AtubB (bottom) superimposed on PDB 7QUP α tubulin (2227 out of 2653 residues superimposed in PyMOL, RMSD = 0.918 Å). 7QUP is grey, AtubAB is green. See also Movie M2.

**Figure S3G.**
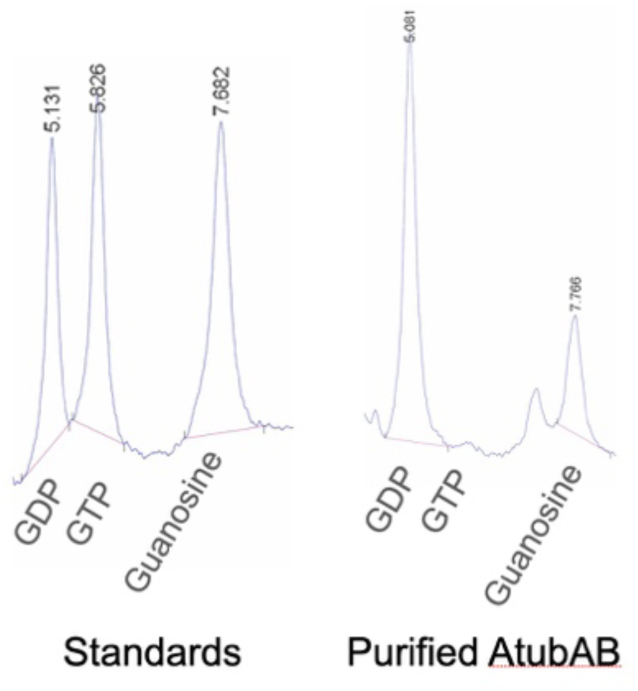
Purified AtubAB mostly contains GDP. The purified untagged AtubAB protein was extracted for nucleotides and using HPLC they were compared against GDP/GTP and guanosine standards (left). The sample on the right shows that purified AtubAB contains very little GTP, meaning that the stable AtubAB heterodimer as purified here (using harsh conditions) does most likely trap a GDP in AtubB, which is in line with the finding that the polymerised protein filaments also contained only GDP (Figure 3K). Numbers refer to elution time in min.

**Figure S3H.**
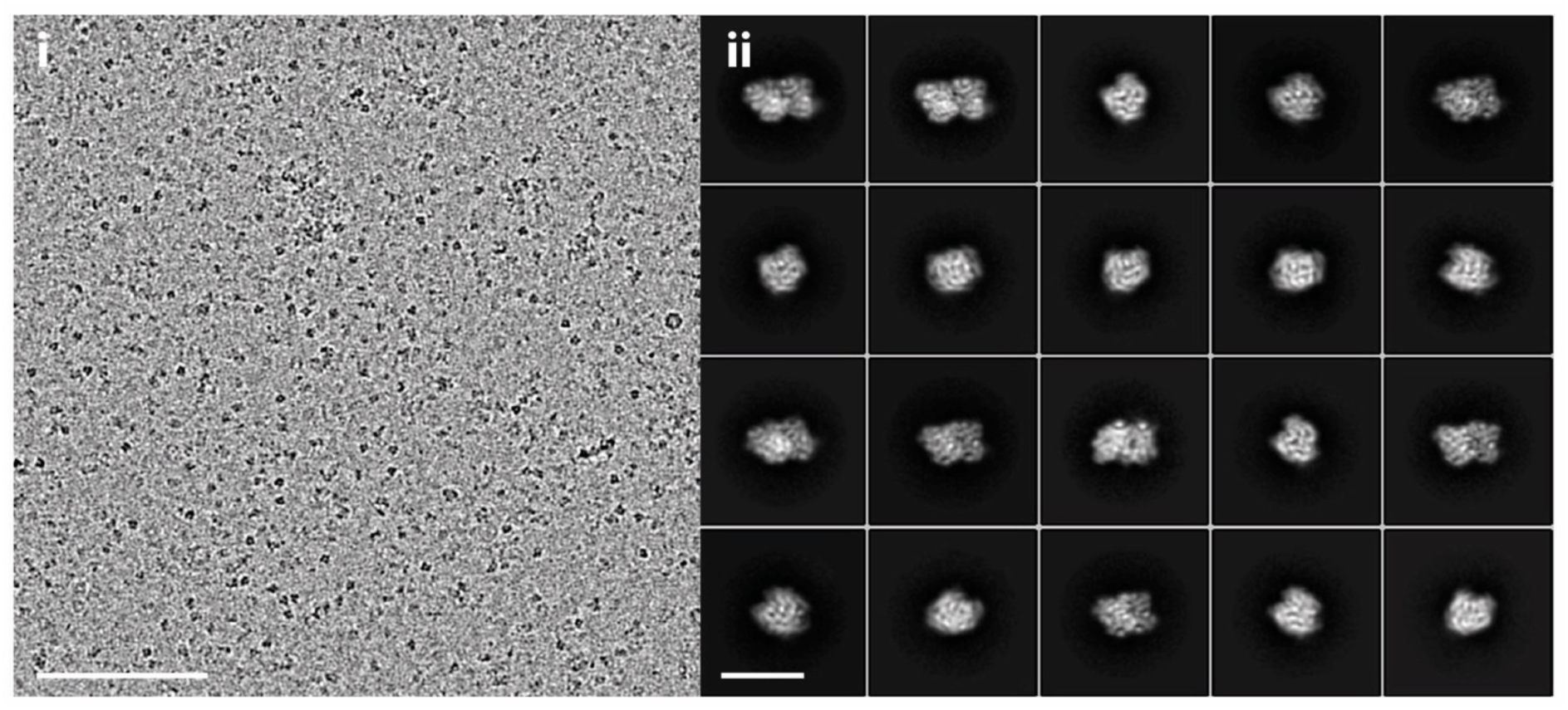
AtubAB cryo-EM for single-particle analysis (SPA). i) cryo-EM image of untagged AtubAB for single particle analysis (SPA; no nucleotide added, structure Figure 3K, right). Bandpass filtered for clarity. Scale bar 100 nm. ii) 2D class averages after picking particles, revealing top and side views of the AtubAB dimers. Scale bar 10 nm.

**Figure S3I.**
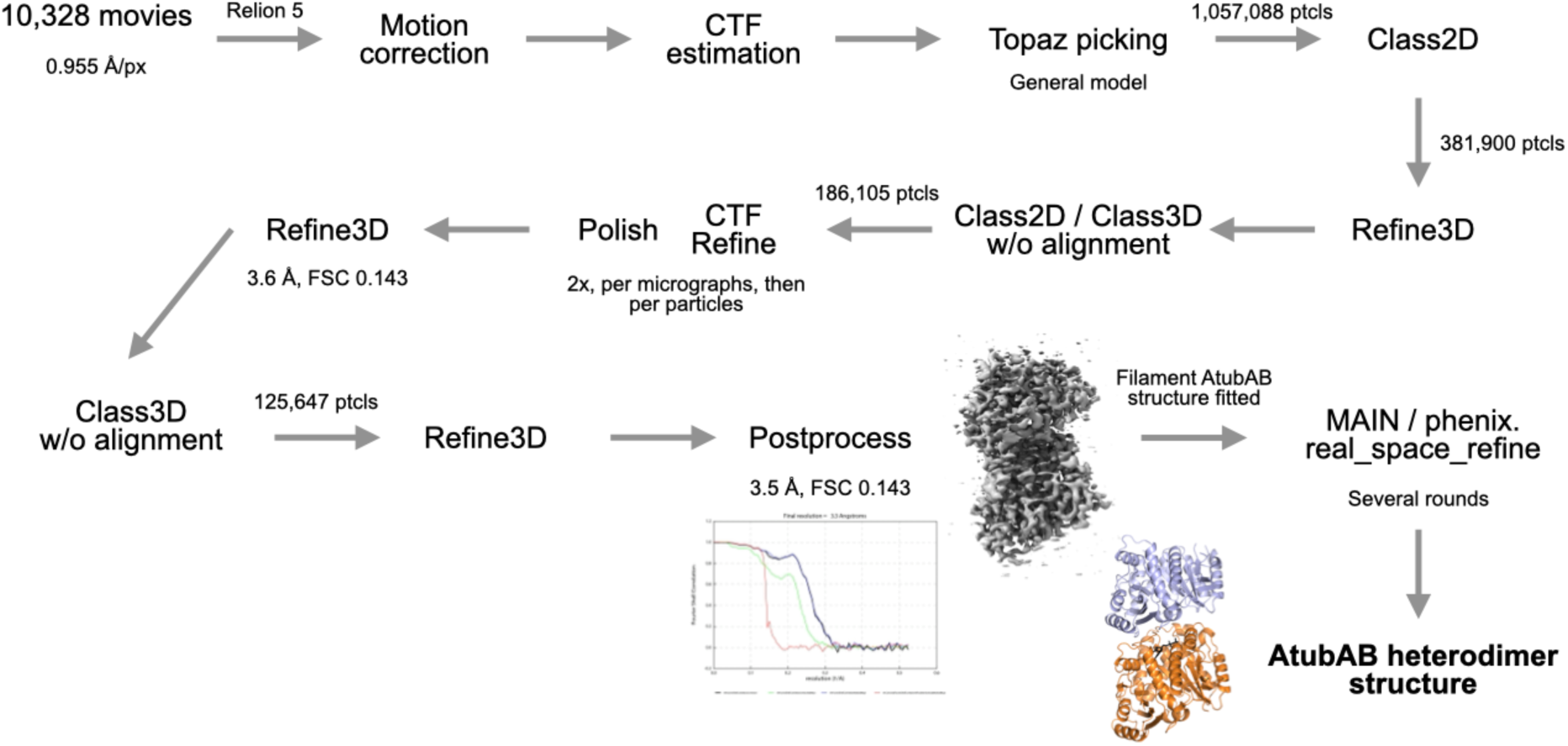
Cryo-EM single-particle processing scheme of untagged AtubAB un-polymerised heterodimer (SPA).

**Figure S4A.**
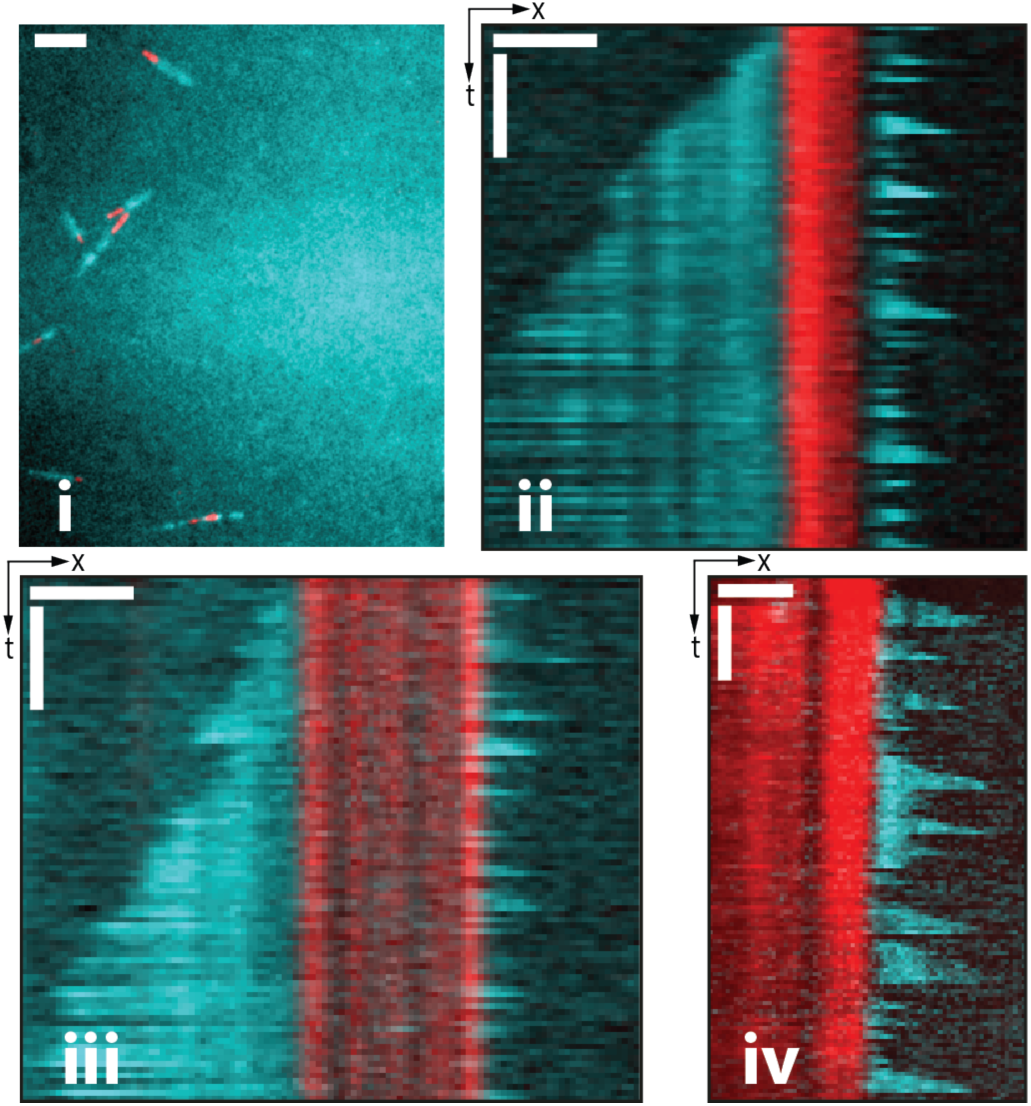
TIRF microscopy of AtubAB filament dynamics. TIRF microscopy, AtubAB polymerisation and kymographs as in Figure 4A-C. Red: GMPCPP-stabilised seeds (not depolymerising), cyan: AtubAB with GTP/GDP, dynamic**. i)** Field of view using untagged proteins, only (4 µM final concentration). Scale bar 5 µm. **ii) and iii)** Further examples of dynamic untagged AtubAB filaments (4 µM final concentration), similar to Figure 4C, showing slow growth at the minus ends (left) and fast growth and dynamic instabilities at the plus ends (right). **iv)** Same as ii and iii for a sample with 20% tagged, unlabelled AtubAhBs added (Methods), to increase turnover at higher concentrations needed to get highly populated field views (as in Figure 4A) (5 µM, see Methods). Scale bars = 2 min and 2 µm.

**Figure S4B.**
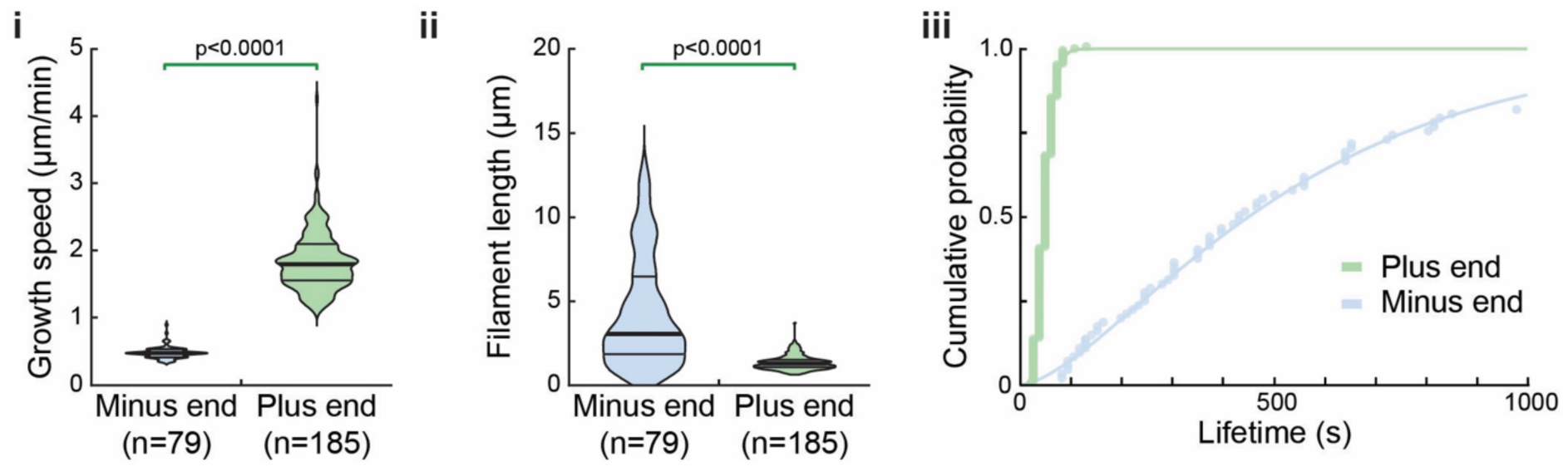
Quantifications of AtubAB mini microtubule dynamics as observed by TIRF microscopy. Relate to Figures 4C and S4A. **i) & ii)** AtubAB GMPCPP seeds were polymerised and filament growth rates **(i)** (minus ends, median speed of growth 0.48 µm/min; plus ends, median speed of growth 1.80 µm/min) and maximal lengths before undergoing catastrophe **(ii)** were measured. p values from two-sided Kruskal-Wallis tests for non-parametric samples are indicated; n, number of individual growth events quantified from at least three independent experiments. Thick lines, median; thin lines, quartile. **iii)** Cumulative AtubAB filament lifetime distributions of mini microtubules grown at the minus (blue) and plus ends (green). Mean lifetime estimates +/- error (lifetime at half cumulative distribution): 451.9 +/- 55.5 s (minus ends) and 45.1 +/- 1.6 s (plus ends). Line, gamma distribution fits.

### SUPPLEMENTARY TABLE

**Supplementary Table 1.** Cryo-EM/ET data, refinement and validation statistics.

|  |  |  |  |
| --- | --- | --- | --- |
|  | AtubAB mini microtubule in *E. coli* cells by cryo-ET with subtomogram averaging; EMD-56154 | AtubAB mini microtubule by cryo-EM; EMD-56153; PDB ID 9TQK | AtubAB free heterodimer by cryo-EM; EMD-56152; PDB ID 9TQJ |
| Proteins | Untagged AtubAB: GenBank MDH5401500.1 (M1 replaced with MSEVVVV); MDH5401501.1 | Untagged AtubAB: GenBank MDH5401500.1 (M1 replaced with MSEVVVV); MDH5401501.1 | Untagged AtubAB: GenBank MDH5401500.1 (M1 replaced with MSEVVVV); MDH5401501.1 |
| **Data and processing** |  |  |  |
| Magnification | 83,000 | 105,000x | 130,000x |
| Voltage (kV) | 300 | 300 | 300 |
| Electron exposure (e-/Å<sup>2</sup>) | 152 e-/Å<sup>2</sup>, 37 tilts at 4.1 e-/Å<sup>2</sup> | 60 | 50 |
| Defocus range (µm) | -1.5 µm to -3.5 µm | -1.5 to -2.5 | -1.5 to -2.5 |
| Pixel size (Å) | 1.514 | 1.222 | 0.955 |
| Symmetry imposed | Twist = -89.824°, rise = 10.248 Å | Twist= -89.836°, rise = 10.460 Å | None |
| Initial particle images (no.) | 47 tomograms, 44 used, 23,734 stacks | 61,534 | 1,057,088 |
| Final particle images (no.) | 15,362 stacks | 27,702 | 125,647 |
| Map resolution (Å) | 6.4 | 3.1 | 3.5 |
| FSC threshold | 0.143 | 0.143 | 0.143 |
| **Refinement** |  |  |  |
| Initial model used (PDB code) | None | *De novo* (Relion 5 - ModelAngelo) | AlphaFold3 model |
| Map sharpening *B* factor (Å<sup>2</sup>) | -200 | -110 | -139 |
| <u>Model composition</u> | Not applicable | AtubAB (heterodimer) | AtubAB heterodimer |
| Non-hydrogen atoms |  | 6543 | 6016 |
| Protein residues |  | A: 418; B: 422 | A: 354; B: 420 |
| Ligands |  | 2 GDP (in A and B) | 1 GDP in AtubB |
| B factors protein (Å<sup>2</sup>) |  | 66.4 | 72.8 |
| RMSD bond lengths (Å) |  | 0.003 | 0.007 |
| RMSD bond angles (°) |  | 0.618 | 1.144 |
| <u>Validation</u> |  |  |  |
| MolProbity score |  | 1.4 (97<sup>th</sup> percentile) | 1.5 (96<sup>th</sup> percentile) |
| Clashscore |  | 6.55 | 8.77 |
| Poor rotamers (%) |  | 0.7 | 0.3 |
| Rama Favoured (%) |  | 97.7 | 98.2 |
| Rama Disallowed (%) |  | 0.0 | 0.0 |

### SUPPLEMENTARY MOVIES

**Movie M1.** Electron cryotomogram (cryo-ET) of FIB-milled lamella of *E. coli* C41(DE3) cell over-expressing untagged AtubAB, showing bundles of mini microtubules crossing a cell division site (same data as shown in Figure 3A, cells expressing AtubAB as shown in Figure 2A, left).

**Movie M2.** Superposition of AtubAB filament dimer structure and eukaryotic (*D. melanogaster*) α/β tubulin in microtubule (PDB 7QUP). 7QUP is grey, AtubAB is green. Same as Supplementary Figure S3F, right.

**Movie M3.** Overview of the architecture of AtubAB four-stranded mini microtubules. Same structure as in Figure 3F.

**Movie M4.** AtubAB’s cytomotive switch, the polymerisation-dependent conformational switch enabling filament dynamics. Morph between AtubB in the filament and AtubB in the un-polymerised dimer conformations (same as Figure 3L).

**Movie M5.** TIRF microscopy of AtubAB filaments revealing dynamic instability. Red: stable GMPCPP seeds. Cyan: dynamic AtubAB (GTP/GDP). The kymographs on the right show two different filaments from the overview on the left. They grow slowly at the minus ends (left side of the red GMPCPP seeds) and grow fast at the plus ends (right side), where they also undergo frequent catastrophes to display dynamic instability (very rapid depolymerisation/shortening). Some data the same as in Figures 4B & C. Scale bar 5 µm.

### SUPPLEMENTARY DATA

FASTA, tree and alignment data will be made available online via Figshare.

### SUPPLEMENTARY DISCUSSION

#### Phylogenetic analyses

Reconstructing reliable phylogenetic trees from tubulin protein sequences is particularly challenging because the alignments contain limited phylogenetic signal. Tubulins are highly conserved, (relatively) short proteins, resulting in low effective sequence variation relative to the number of taxa analysed. This low signal amplifies the impact of stochastic errors, model misspecification, and alignment uncertainty on inferred topologies. Consequently, deep or rapid divergence events are especially difficult to resolve, and alternative tree topologies often receive comparable statistical support. In this case, this translates to multiple clades robustly inferred as monophyletic (eukaryotic tubulin α, β, γ, δ, ε and cryptic clades, AtubA + BtubA, AtubB + BtubB, AtubC, and Odin tubulin), but with unclear relatedness patterns between them.

To identify recurrent topological patterns that may illuminate the evolution of tubulin sequences, we performed additional phylogenetic reconstructions varying sequence sampling. To assess possible effects of long-branch attraction due to long-branching outgroup sequences, we performed phylogenetic analyses including only the ingroup (that is, eukaryotic and prokaryotic tubulins only), or including combinations of three tubulin homologues: artubulin, CetZ, and a small group of Asgard archaeal sequences that may be associated to CetZ.

The results from these phylogenies corroborated the main findings in Figure 1A, including: (1) BtubA and BtubB clustering within AtubA and AtubB; (2) AtubA and AtubB clustering with eukaryotic α tubulin or at the base of the α and β tubulin clade; and (3) AtubC clustering with Odin tubulin (Supplementary Discussion Figures 1 & 2, below). While the affiliation of AtubC and Odin tubulin in the tubulin clade was not always clear, they tended to branch as part of a monophyletic group with α and β tubulins, AtubA-BtubA, AtubB-BtubB, and, sometimes, γ tubulin. This result remained consistent after trimming the sites with the lowest decile of low-likelihood scores (option “--robust-phy 0.9” in IQ-Tree v3.0.1) (Supplementary Discussion Figure 3) (Liu et al., 2025).

Notably, as explained above, the obtained results lack sufficient phylogenetic signal to produce robust branch support values. For this reason, we employed the Transfer Bootstrap Expectation (TBE) metric (Lemoine et al., 2018), which considers the average number of sequences that are ‘transferred’ across the bipartition in bootstrap trees, compared to the number of leaves at the light side of the bipartition (i.e. a TBE value of 0.9 indicates that 10% of the number of leaves in the light side of the bipartition are transferred on average). Thus, when applied to clades including groups with very few members, high support values need to be considered with a high degree of caution.

Given the very long branches present in the tubulin ε δ and cryptic clades, we considered that their position at the base of the tubulin tree may be the result of long-branch attraction. We therefore reconstructed trees after removing these groups to evaluate whether AtubABC and/or Odin tubulins would then cluster at the base. These trees indicated Odin tubulins as the first group to diverge, followed by AtubC, regardless of whether the outgroup was defined by CetZ, artubulin, or both (Supplementary Discussion Figures 4 - 6).

Altogether, we were unable to determine, with very high confidence, a specific evolutionary trajectory for Asgard archaeal and eukaryotic tubulins. While histories involving horizontal transfers are consistent with these results, given the multiple branching patterns of Asgard archaeal tubulins in the tubulin tree and following the long series of recent studies finding an Asgard archaeal ancestry of eukaryotic proteins, we favour a scenario in which duplications and losses were common for tubulin homologues in Asgard archaea, a subset of which was inherited by the eukaryotic lineage, as is discussed in the main text.

## Supplementary Discussion Figures

**Supplementary Discussion Figure 1.**
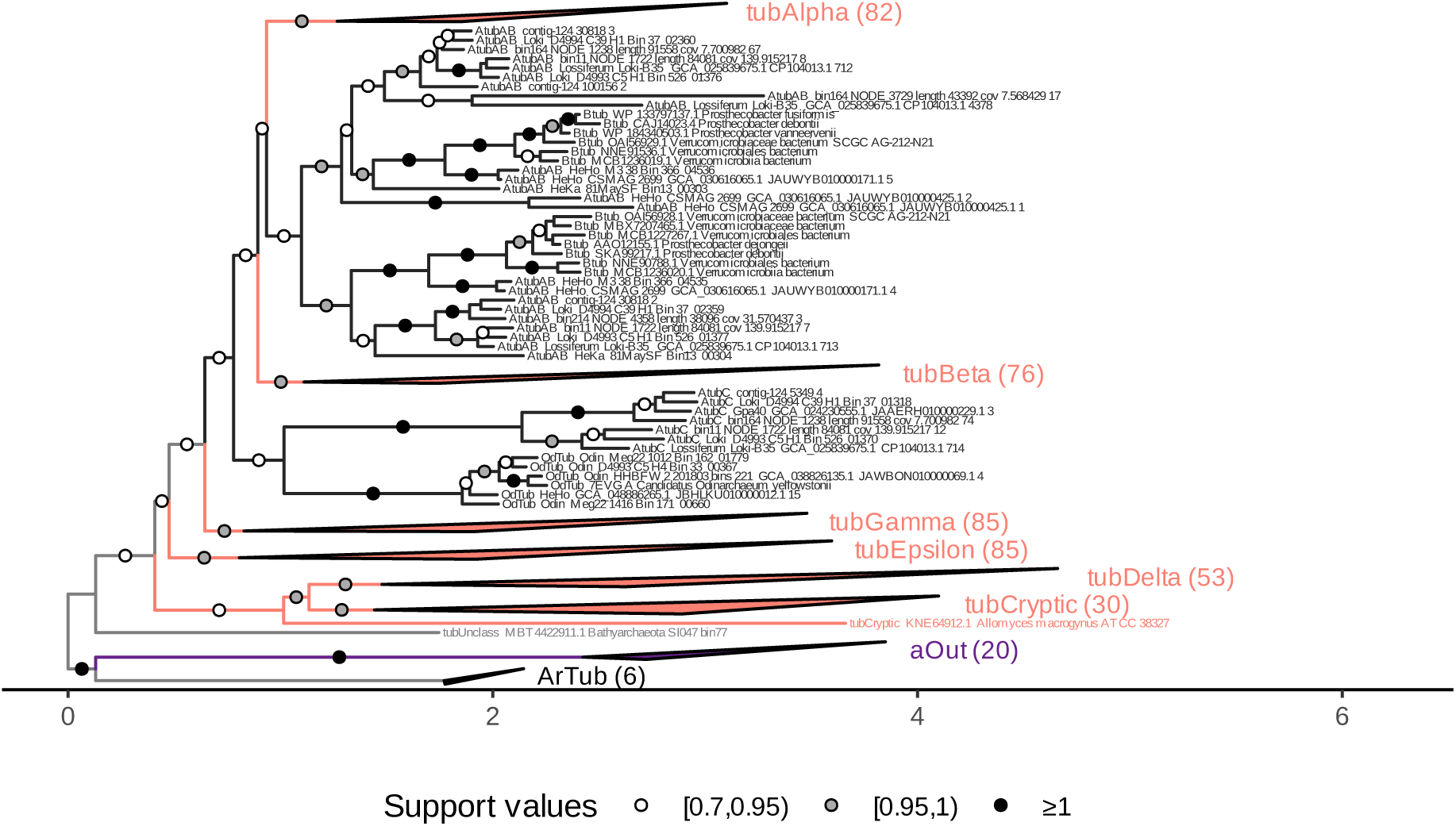
Phylogeny reconstructed using IQ-TREE 3 under the model Q.pfam+C50+G4+PMSF of an alignment including artubulin and a CetZ-related Asgard archaeal clade as outgroup, aligned with MAFFT-linsi, trimmed with trimAl of sites containing over 50 % gaps, pruned of sequences formed by over 50 % gaps, containing 488 sequences and 431 columns. Branch support values represent Transfer Bootstrap Expectation. Branch length legend indicates substitutions per site. The same tree, indicating Felsenstein Bootstrap Proportions is shown in Supplementary Discussion Figure 7.

**Supplementary Discussion Figure 2.**
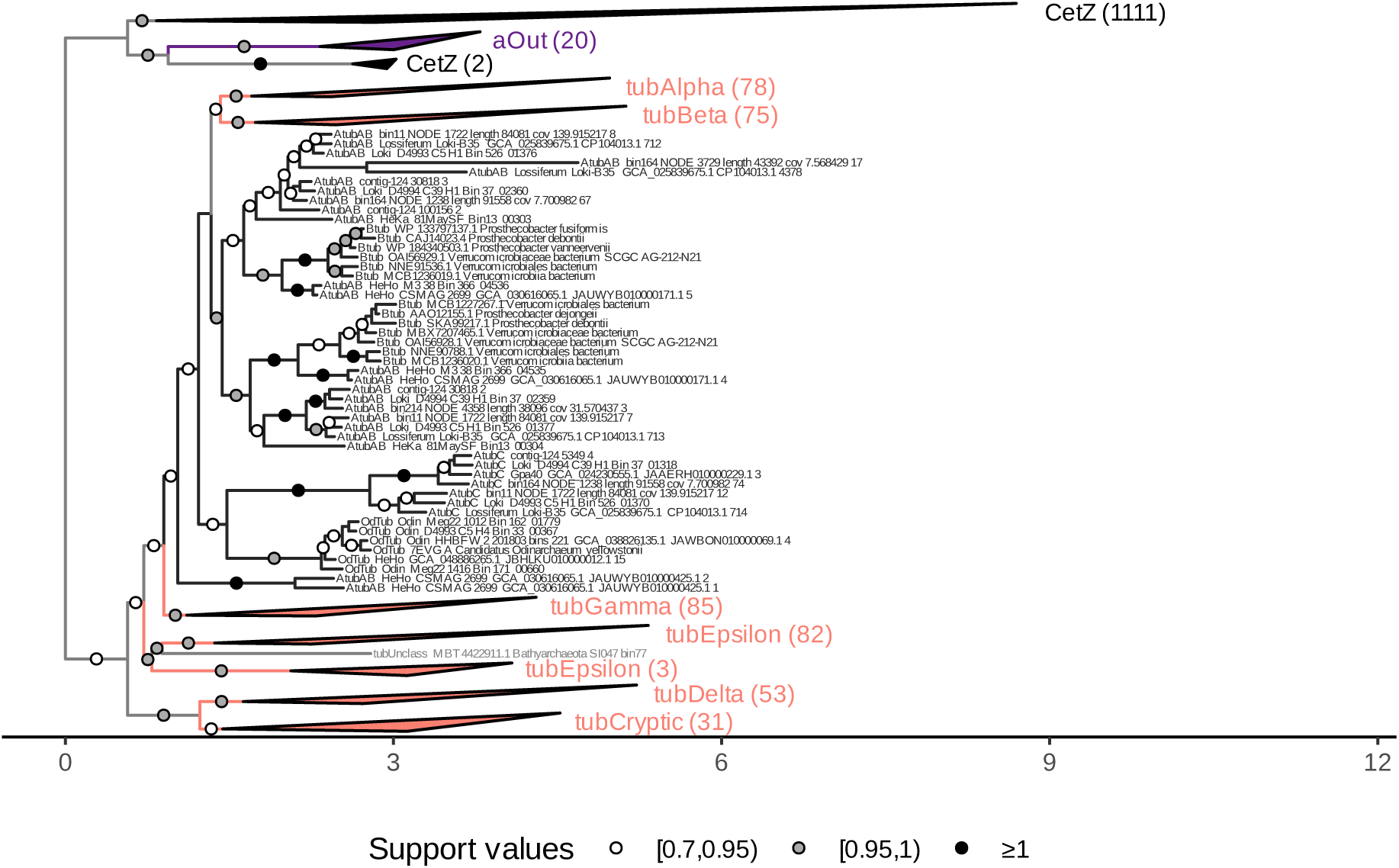
Phylogeny reconstructed using IQ-TREE 3 under the model Q.pfam+C50+G4+PMSF of an alignment including CetZ and a CetZ-related Asgard archaeal clade as outgroup, aligned with MAFFT-linsi, trimmed with trimAl of sites containing over 50 % gaps, pruned of sequences formed by over 50 % gaps, containing 1590 sequences and 337 columns. Branch support values represent Transfer Bootstrap Expectation. Branch length legend indicates substitutions per site. The full tree, indicating Felsenstein Bootstrap Proportions is shown in Supplementary Discussion Figure 8.

**Supplementary Discussion Figure 3.**
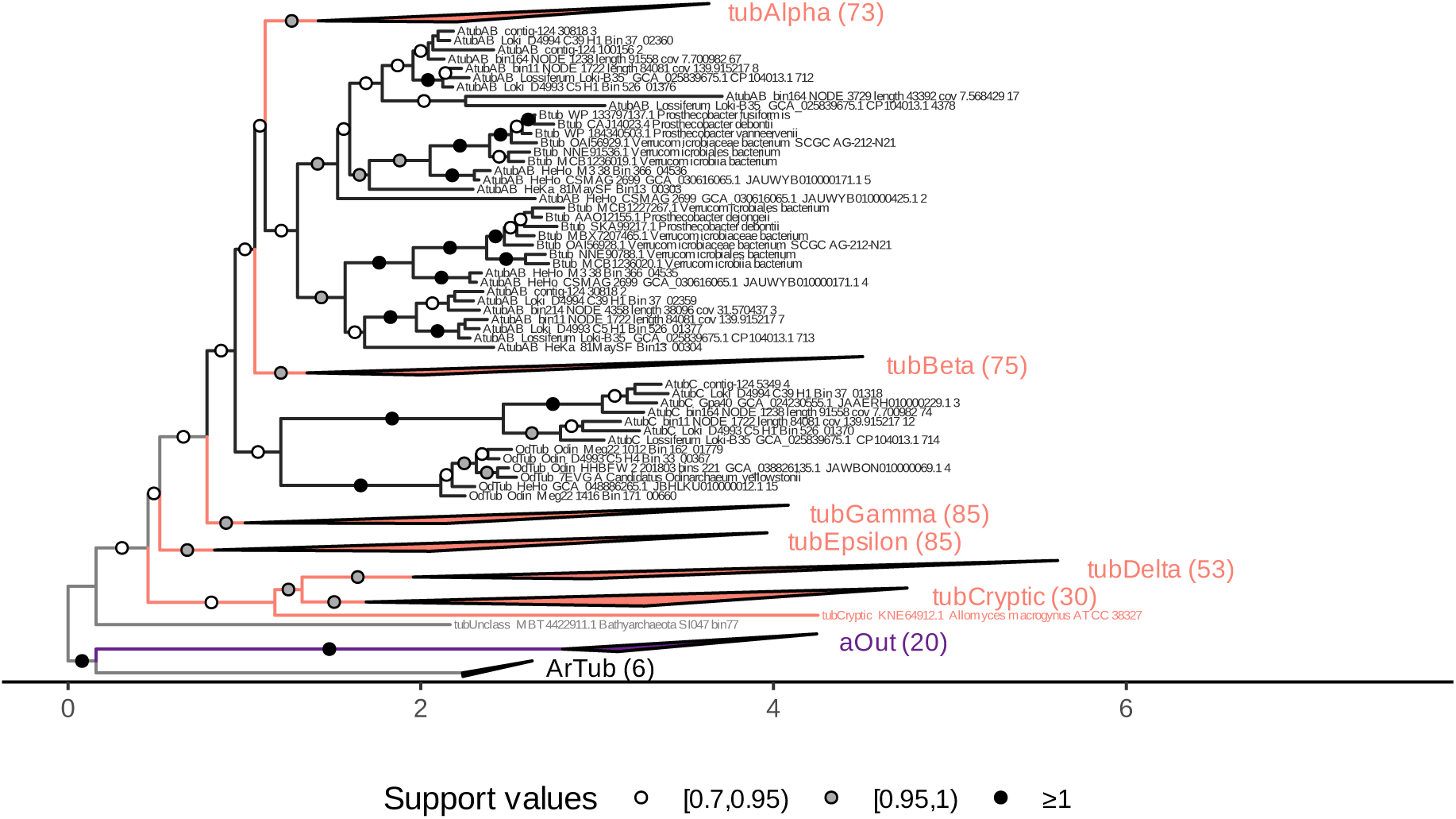
Phylogeny reconstructed using IQ-TREE 3 under the model Q.pfam+C50+G4+PMSF of an alignment including artubulin and a CetZ-related Asgard archaeal clade as outgroup, aligned with MAFFT-linsi, trimmed with trimAl of sites containing over 90 % gaps, pruned of sequences formed by over 50 % gaps, and reconstructed while trimming the 10 % of sites with the lowest likelihood scores during the reconstruction with IQ-TREE 3. Tree used as input to IQ-TREE 3 contained 477 sequences and 578 columns. Branch support values represent Transfer Bootstrap Expectation. Branch length legend indicates substitutions per site. The full tree, indicating Felsenstein Bootstrap Proportions is shown in Supplementary Discussion Figure 9.

**Supplementary Discussion Figure 4.**
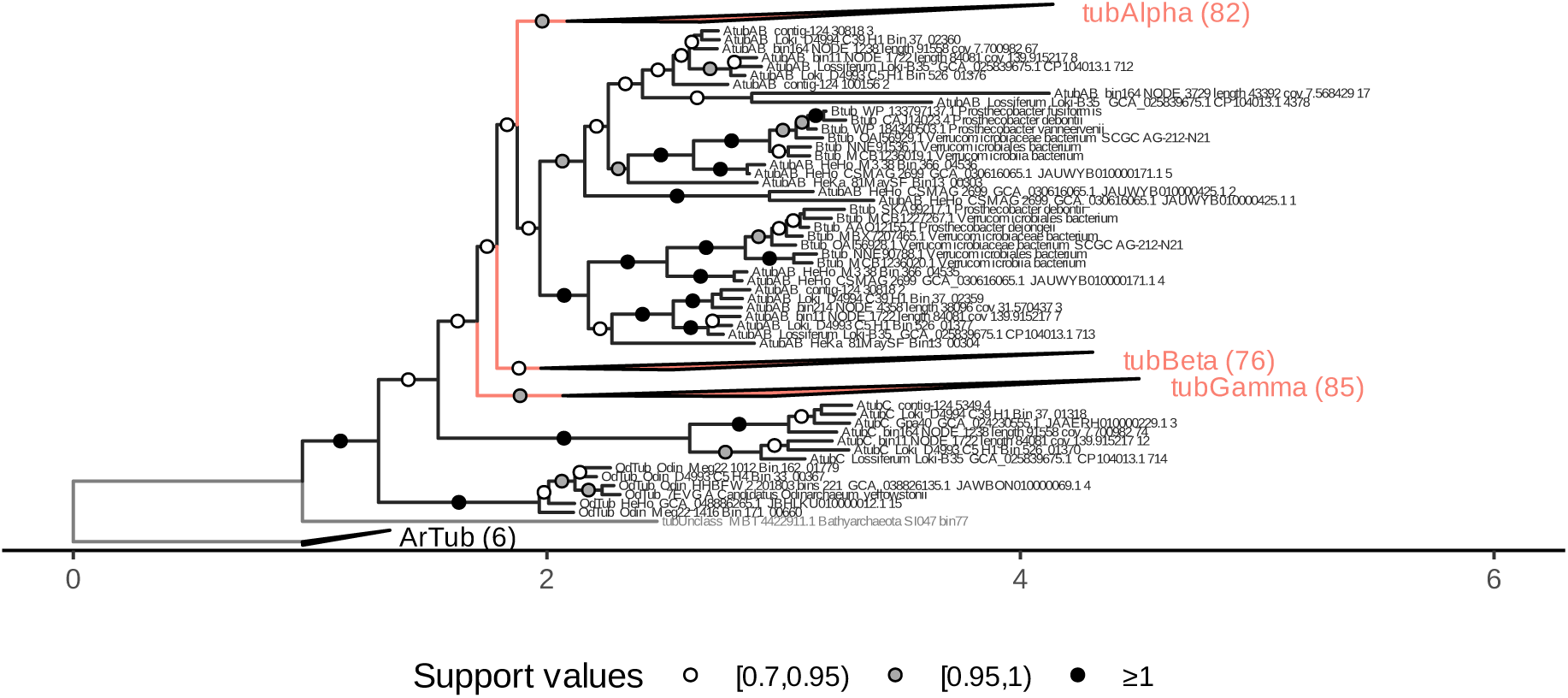
Phylogeny reconstructed using IQ-TREE 3 under the model Q.pfam+C50+G4+PMSF of an alignment including artubulin as outgroup, aligned with MAFFT-linsi, trimmed with trimAl of sites containing over 50 % gaps, pruned of sequences formed by over 50 % gaps, and reconstructed while trimming the 10 % of sites with the lowest likelihood scores during the reconstruction with IQ-TREE 3. Tree used as input to IQ-TREE 3 contained 299 sequences and 435 columns. Branch support values represent Transfer Bootstrap Expectation. Branch length legend indicates substitutions per site. The full tree, indicating Felsenstein Bootstrap Proportions is shown in Supplementary Discussion Figure 10.

**Supplementary Discussion Figure 5.**
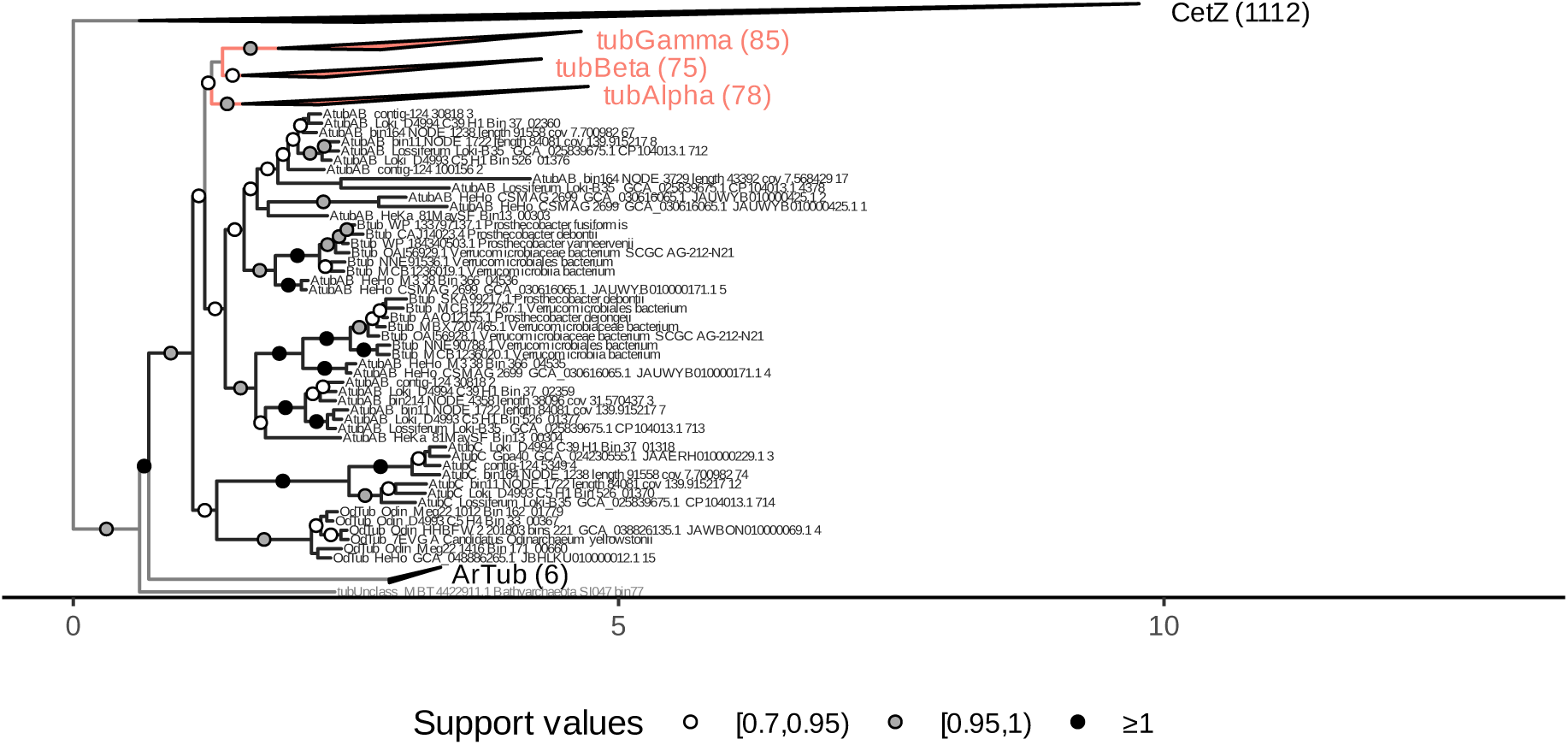
Phylogeny reconstructed using IQ-TREE 3 under the model Q.pfam+C50+G4+PMSF of an alignment including artubulin and CetZ as outgroup, aligned with MAFFT-linsi, trimmed with trimAl of sites containing over 50 % gaps, pruned of sequences formed by over 50 % gaps, containing 1406 sequences and 339 columns. Branch support values represent Transfer Bootstrap Expectation. Branch length legend indicates substitutions per site. The full tree, indicating Felsenstein Bootstrap Proportions is shown in Supplementary Discussion Figure 11.

**Supplementary Discussion Figure 6.**
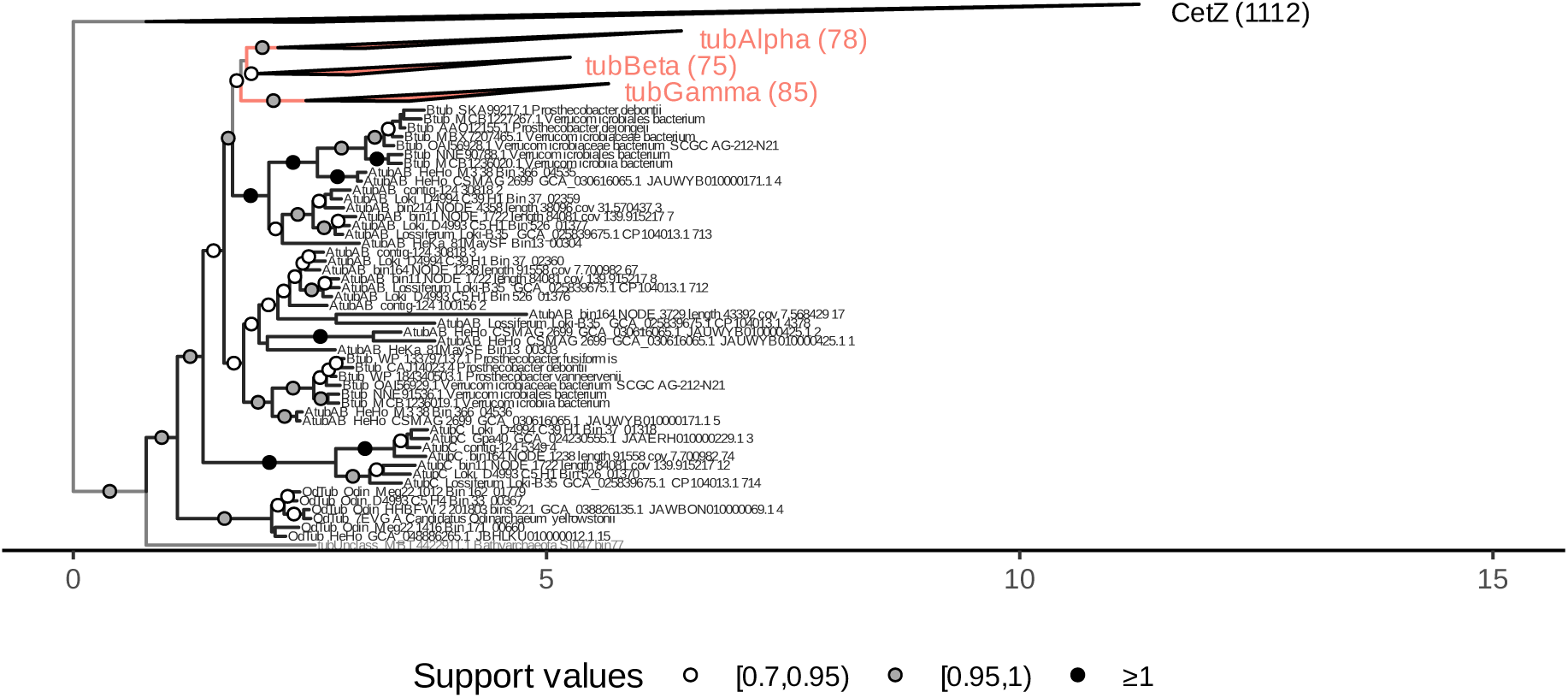
Phylogeny reconstructed using IQ-TREE 3 under the model Q.pfam+C50+G4+PMSF of an alignment including CetZ as outgroup, aligned with MAFFT-linsi, trimmed with trimAl of sites containing over 50 % gaps, pruned of sequences formed by over 50 % gaps, and reconstructed while trimming the 10 % of sites with the lowest likelihood scores during the reconstruction with IQ-TREE 3. Tree used as input to IQ-TREE 3 contained 1400 sequences and 337 columns. Branch support values represent Transfer Bootstrap Expectation. Branch length legend indicates substitutions per site. The full tree, indicating Felsenstein Bootstrap Proportions is shown in Supplementary Discussion Figure 12.

**Supplementary Discussion Figure 7.**
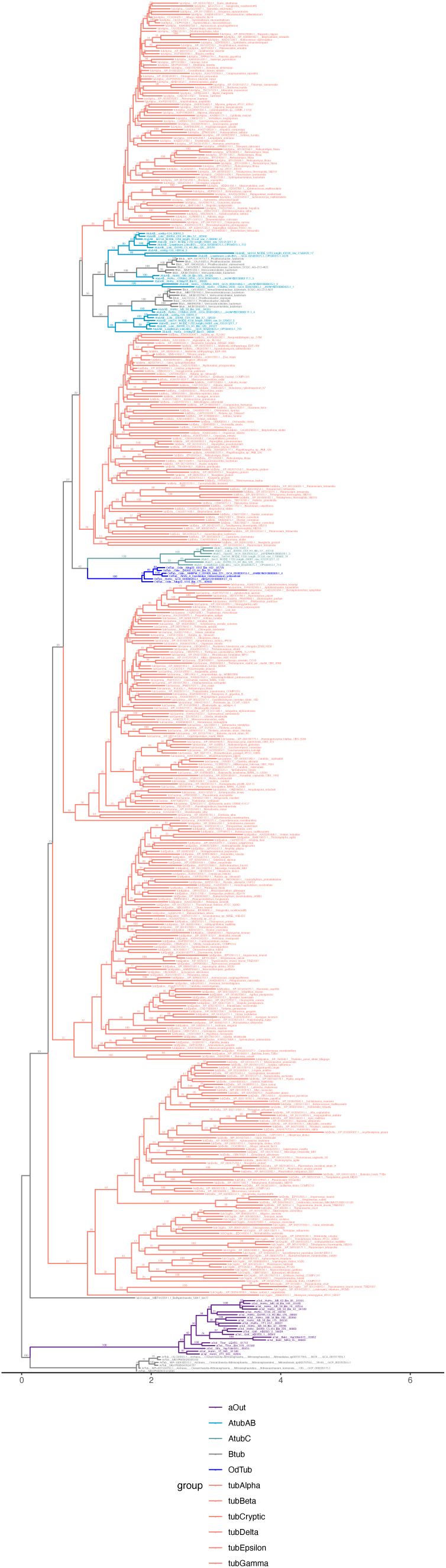
Full phylogeny corresponding to Supplementary Discussion Figure 1, indicating Felsenstein Bootstrap Proportions as branch support values.

**Supplementary Discussion Figure 8.**
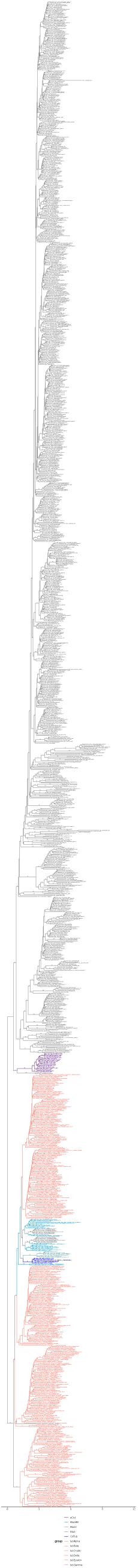
Full phylogeny corresponding to Supplementary Discussion Figure 2, indicating Felsenstein Bootstrap Proportions as branch support values.

**Supplementary Discussion Figure 9.**
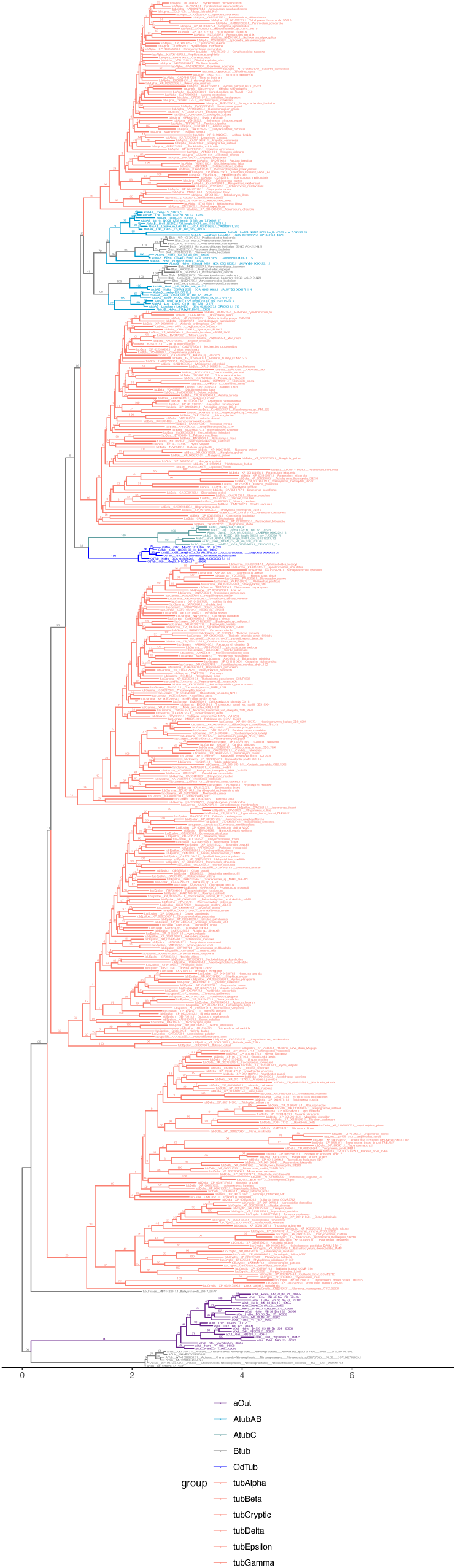
Full phylogeny corresponding to Supplementary Discussion Figure 3, indicating Felsenstein Bootstrap Proportions as branch support values.

**Supplementary Discussion Figure 10.**
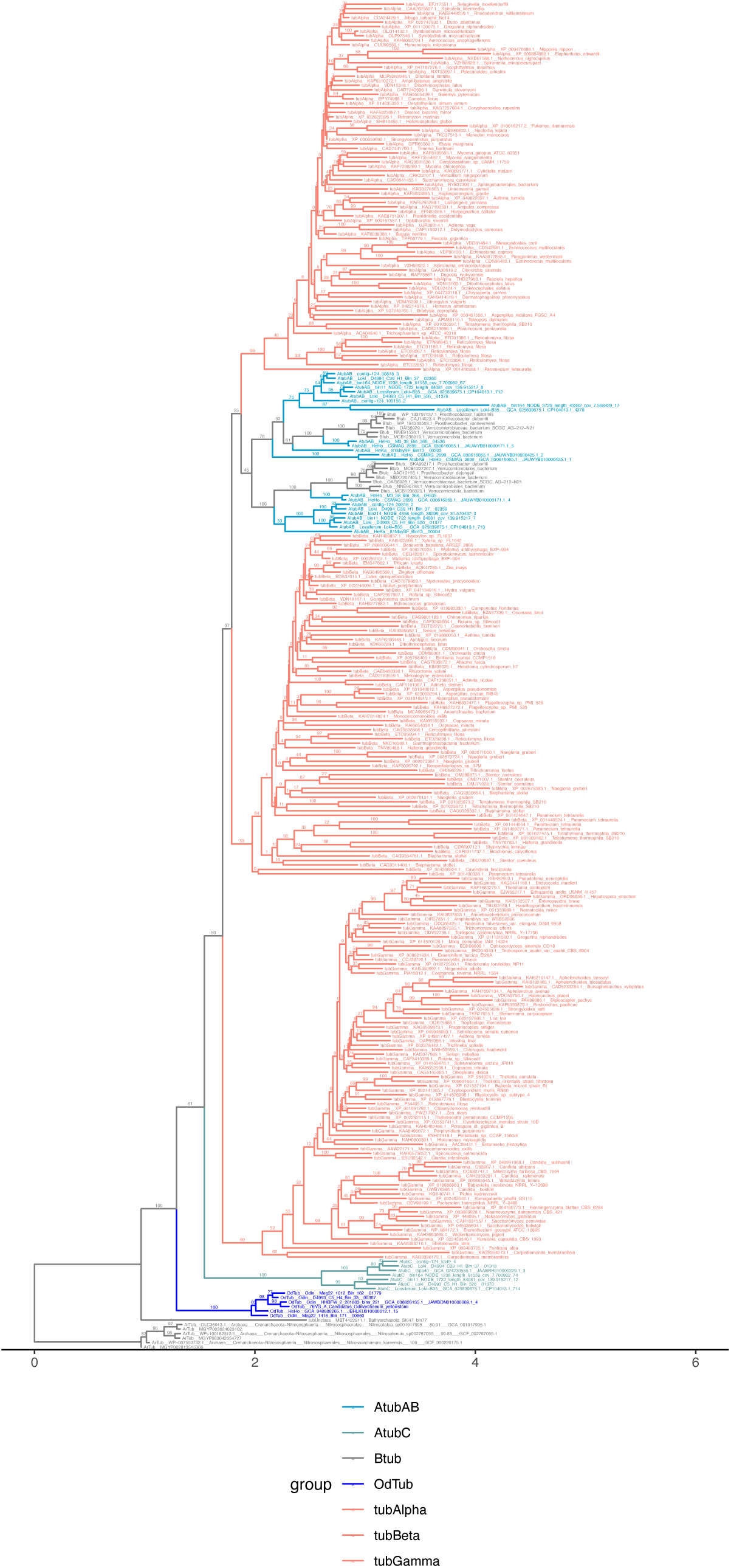
Full phylogeny corresponding to Supplementary Discussion Figure 4, indicating Felsenstein Bootstrap Proportions as branch support values.

**Supplementary Discussion Figure 11.**
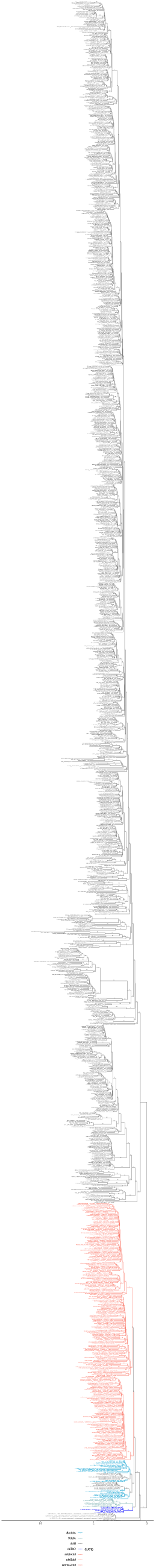
Full phylogeny corresponding to Supplementary Discussion Figure 5, indicating Felsenstein Bootstrap Proportions as branch support values.

**Supplementary Discussion Figure 12.**
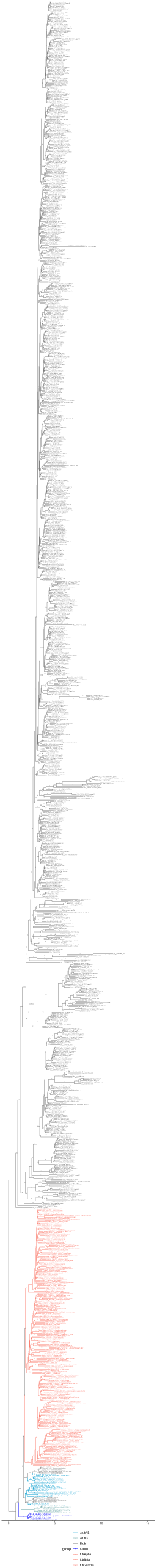
Full phylogeny corresponding to Supplementary Discussion Figure 6, indicating Felsenstein Bootstrap Proportions as branch support values.

